# A complex adaptive systems model of labor reciprocity and normative reasoning in swidden agriculture

**DOI:** 10.1101/2024.12.12.628215

**Authors:** Denis Tverskoi, Shane A. Scaggs, Sean S. Downey

## Abstract

Swidden (aka “slash-and-burn”) agriculture is a prototypical coupled human and natural system and understanding the social, cultural, and environmental factors shaping the trajectories of swidden forests is essential to mitigate climate change and ensure equitable collaboration between scientists, planners, and Indigenous communities. Despite this, mathematical models integrating its social and ecological dynamics are rare. Here, we use complex adaptive systems theory to develop a model where individuals rely on labor exchange driven by reciprocity, and on normative reasoning that can lead to sanctions. Our results identify three emergent regimes: low-intensity swidden, sustainable high-intensity swidden that maximizes ecosystem services and harvest returns, and deforestation. We show that sustainable high-intensity swidden evolves if labor reciprocity and normative reasoning are balanced: helping behavior should be significantly conditioned by normative reasoning to prevent over-harvesting, while reciprocity is necessary to prevent excessive sanctioning. We find that the sustainable high-intensity swidden regime is robust to changes in group size, is resilient to environmental shocks, can evolve under various models of forest ecology, and is most productive for both forests and farmers when the balance of labor reciprocity and normative reasoning results in an intermediate scale of forest disturbance. Overall, we illuminate the importance of Indigenous social norms and customary practices related to swidden labor in maintaining sustainable and intensive swidden agriculture.

## 1 Introduction

Swidden involves the intentional use of fire to clear primary or secondary forests to create agricultural fields (Jiang *et al*., 2022). Fire eliminates pests and release organic matter and nutrients bound in plant biomass into the topsoil. In the swidden farming cycle, short-term cultivation is typically followed by relatively long periods of fallowing (Dove and Kammen, 1997; Jiang *et al*., 2022). This type of farming has been practiced for thousands of years (Dumond, 1961; Bellwood *et al*., 2007; Ribeiro Filho *et al*., 2013) and is now widely used in Indigenous and smallholder communities in tropical and subtropical regions, including Central and South America, South Asia, and Africa (Downey, 2009; Jakovac *et al*., 2016). Estimates suggest that between 300 million (Dove, 1983) and 1 billion people (Ribeiro Filho *et al*., 2013) rely on swidden agriculture to some extent. In 2018, Indigenous peoples controlled 37% of the natural land on Earth (Garnett *et al*., 2018). 36% of intact forests (Fa *et al*., 2020), 60% of all well-known terrestrial mammals (O’Bryan *et al*., 2021), and 24% of the total above-ground carbon in the world’s tropical forests (Fund and Center, 2015) are on Indigenous peoples’ lands. These estimates indicate that swidden agriculture, along with other traditional forms of forest-based subsistence, significantly affect global climate dynamics and sustainability. As Pisor et al., and others demonstrated in a recent special issue, “Climate change adaptation needs a science of culture” (Pisor *et al*., 2023). Therefore, understanding and acknowledging the role of Indigenous swidden practices in biodiversity conservation and forest stewardship is crucial (Fernández-Llamazares *et al*., 2024).

The literature on swidden is vast and has been reviewed elsewhere (e.g., Cairns, 2015), however we provide a brief overview here. Historically, perspectives on the effects of swidden agriculture on the environment and the conservation of tropical forest ecosystems have been polarized. Some research emphasizes its negative effects on the environment, hypothesizing that it typically leads to soil degradation and deforestation – especially in the context of population growth – and advocates for the “modernization” of traditional agricultural practices (see Ribeiro Filho *et al*. (2013) for examples). A substantial amount of research relates to swidden farming practices such as the length of the fallow period, managing crop diversity, land preparation methods, and traditional ecological knowledge (Eden and Andrade, 1987; Kleinman *et al*., 1995; Mertz *et al*., 2009; Ford and Nigh, 2016). An increasing number of researchers are also using an integrated socioecological system approach in which human social, cultural and demographic factors co-evolve with tropical forest ecosystem dynamics (Dove, 1983; Rappaport, 2000; Barton, 2014; Downey, 2010). This approach typically views swidden agriculture as a productive and sustainable system that can under some circumstances enhance the forest biodiversity (Pedroso-Junior *et al*., 2009; Padoch and Pinedo-Vasquez, 2010; Ford and Nigh, 2016; Downey *et al*., 2023; Balée, 2023). A smaller body of research examines how social factors such as social structure, social norms, religion, and rituals may also be important determinants of sustainable swidden cultivation (Downey *et al*., 2020; Rappaport, 2000). Indeed, swidden farming typically involves numerous labor-intensive tasks such as forest clearing, burning, planting, weeding and harvesting, and many of these activities can be performed more safely and efficiently when people work together in groups. As a result, both social and environmental factors contribute to the dynamics of labor exchange networks and may also shape the trajectory and outcome of local cultivation practices. Clearly, a deep understanding of both the social and environmental drivers of sustainable swidden agriculture is crucial for the conservation of tropical ecosystems and the mitigation of climate change.

Long-term ethnographic research in Maya communities that use swidden cultivation in southern Belize indicates that community forests are a common resource, with no formal institution or leader regulating individual cultivation practices (Downey *et al*., 2020). Consequently, swidden labor exchange networks result from social norms related to cooperation rather than institutional leadership. One such norm is direct reciprocity (Roberts, 2008). Direct reciprocity is defined as “a mechanism where people help those who have helped them in the past” (Romano *et al*., 2022). A household survey conducted in two Maya villages in 2018 revealed that direct reciprocity is a significant determinant of the structure of the labor exchange networks associated with swidden cultivation, but that there are also asymmetric labor exchange patterns (Scaggs and Downey, 2024). One possible explanation for this asymmetry is what we define here as *normative reasoning and sanctioning* (or simply, *normative reasoning*) – the cognitive process by which individuals evaluate whether to provide swidden labor to a partner requesting help, based on perceptions, beliefs, or understanding of the acceptability of the request. This decision process was documented in an experimental study involving 150 participants from the same two villages (Downey *et al*., 2020). In the experiment, participants played a common pool resource (CPR) game that simulated swidden cultivation practices in the village. The environment was represented as a grid in which each cell was either occupied by trees or empty, and the rate of forest regeneration increased with the number of forest cells remaining on the grid. Groups of five players simultaneously chose the number of cells they would like to clear, and the game was played over several rounds. In the first experiment, no helpers were required to clear new fields so the behavior of participants in almost all groups converged close to the Nash equilibrium, indicating that participants cleared the maximum possible number of cells in each round, which in turn led to rapid deforestation. In the second experiment, each player was able to clear one cell on their own, but clearing more cells required the help of others, simulating labor exchange needs in the villages. The results demonstrate a significant increase in the remaining fraction of forest among groups. We suspect that this is because participants used normative reasoning when making their helping decisions: information about each player’s requests were displayed to the group, and the results suggested that they used this information to decide whether to provide help, depending on how acceptable they thought the request was. In this way, players could effectively “sanction” (Ostrom, 2002) a player who wanted to clear relatively large fields by refusing to help them, and they could help those who requested smaller fields that they thought were less likely to threaten the common-pool forest resource. Support for this sanctioning mechanism was noted in post-game interviews by one participant: “… some ask for five. If they ask for five, they shouldn’t be helped but if you ask for three, that is a good piece of land and it considers other people”(quoted in Downey *et al*., 2020)^1^. Here we build on these experimental and ethnographic results by employing mathematical modeling to better understand theoretically how normative reasoning and labor reciprocity may shape the evolution of sustainable swidden agriculture.

The existing modeling literature on CPR games is extremely large (Ostrom, 2002; Malézieux and Spiegelman, 2024). Here, we focus on dynamical CPR games with explicit resource dynamics. There are two approaches to modeling such situations. The first approach employs classical game theory assuming that all agents are perfectly rational, so they have the correct beliefs about the behavior of others and the resource dynamics (Antoniadou *et al*., 2013; Noailly *et al*., 2007; Safarzynska, 2018, 2020). Such agents are postulated to maximize their expected discounted utility (payoff) over all future rounds (Antoniadou *et al*., 2013; Safarzynska, 2020). Although this approach is analytically tractable, the assumption of perfect rationality is highly questionable. An alternative approach utilizes evolutionary game theory and is based on the assumption of bounded rationality, implying that agents are not able to fully predict the behavior of others and the dynamics of the environment. Typically, three types of agents are considered: punishers, cooperators, and defector. Cooperators and punishers are prescribed to extract fewer resources than defectors. Punishers sanction defectors, imposing costs on defectors but also being costly for punishers (Sethi and Somanathan, 1996; Noailly *et al*., 2007; Safarzynska, 2013). The within-group dynamics that determine how the frequencies of the three types of agents change are governed by the diffusion of harvesting strategies that generate above-average payoffs, resulting in replicator dynamics. Potential mechanism leading to such dynamics can be natural selection (genetic evolution) where maladapted agents “die” and are removed from a simulation, and payoff-based imitation (cultural evolution) where maladapted agents adopt strategies from agents with higher payoffs. Here, we build a parsimonious agent-based model using a minimalist cultural evolutionary framework. We account for the individual decision-making process by assuming bounded rationality of agents, and we model the social dynamics of swidden labor and a range of simple forest growth dynamics. As a result, we contribute to the literature on modeling dynamical CPR games by considering punishment in the context of labor exchange networks, which is relevant in many natural resource management contexts.

Many existing models of swidden agriculture examine land use and land cover change (LULCC) patterns by linking raster images from remote sensing data with agent-based models of human land use (McCracken, 2002; Ngo *et al*., 2011; Evans *et al*., 2011; Cabrera *et al*., 2012; Assaf *et al*., 2021). Typically, these studies present relationships between demographic variables such as household size, village population levels and land use rates with different levels of agricultural or economic efficiency, and model the decision-making process of agents using assumptions of perfect or bounded rationality (Manson and Evans, 2007). While many existing LULCC swidden models include computationally sophisticated, theoretically informed mechanisms for predicting patterns of land use change in space and time, or the effects of nature conservation policy (Assaf *et al*., 2021), they typically do not incorporate social or cultural factors that can affect land use. A second category of models examines swidden as an integrated socioecological system (SES) with dynamic, integrated social and environmental modeling elements (Baum *et al*., 2016; Guzmán *et al*., 2018; Iwamura *et al*., 2014; Jepsen *et al*., 2006). Most of these models have a large number of free parameters or incorporate raster images from real study locations and therefore they cannot be considered parsimonious models. Nor do they typically account for important social, cultural, and behavioral determinants of swidden agriculture from local communities, with the exception of the demography. In this respect, the model developed by Barton (2014) stands out for its parsimonious design and because it examines settlement patterns related to swidden agriculture and the underlying household, social, and environmental dynamics, including social norms related to land ownership in the ancient American Desert Southwest.

Barton’s model is particularly relevant to our current study because it applies complex adaptive systems (CAS) theory to agent-based modeling and archaeology. CAS theory proposes that most complex systems are characterized by non-linearity and sensitivity to initial conditions, and that low-level interactions among agents can lead to emergent system-level properties (Lansing, 2003). Previously, Downey *et al*. (2023) argued that complex adaptive systems theory is an useful heuristic model for swidden agriculture. The mathematical model we present here follows up on that suggestion. It differs from most of the swidden models reviewed above: it is a parsimonious model that is designed for a comprehensive theoretical exploration of the coupled dynamics of social and behavioral determinants of swidden agriculture. It accounts for social norms related to swidden agriculture, a range of simple ecological models, and emergent properties related to the long-term sustainability of the coupled human and natural system.

Our review of work in evolutionary theory, modeling of CPR games, and computational models of swidden agriculture reveals a gap in our theoretical understanding of the relationship between social norms and customary practices in swidden commons. Here, we hypothesize that labor reciprocity and normative reasoning based on individual perceptions of acceptable land use rates will determine the trajectory and the outcome of the coupled socioecological swidden system. However, it remains unclear (1) how individual perception of acceptability co-evolves with the ecosystem and relates to sustainability; (2) how the interplay of reciprocity and normative reasoning shapes the dynamics of this socioecological system; and (3) what are the joint effects of the above socio-behavioral factors with material, demographic, and ecological factors on the dynamics and sustainability of swidden cultivation. Addressing these issues theoretically is the goal of this paper. To do this, we use a complex adaptive systems framework to develop a mathematical model that focuses on the labor exchange dynamics taking place in groups practicing swidden cultivation in community forests. In doing so, we hope to make the following contributions: first, we contribute to the literature on common pool resource games by modeling the effects of social sanctioning when labor exchange networks are coupled with simple ecological models. Second, we illuminate the importance of social norms and customary practices related to swidden labor in maintaining sustainable and intensive swidden agriculture in community forests worldwide. Third, we document our model in reproducible manner so that other researchers can adapt and integrate it in future research and policy applications.

## 2 The Model

We consider a group of *N* agents interacting on an *L × L* grid over *T* rounds. We treat this group as a village and each agent as a household.

### Ecological dynamics

Following Bak *et al*. (1990), we postulate each cell *k* is either occupied by trees or empty. Note that we intentionally simplify the ecological dynamics and do not model the different stages of forest succession, because our focus is on the social dynamics shaping the evolution of sustainable swidden agriculture (see Downey *et al*., 2023, for an analysis of succession in swidden forests). We assume that each empty cell turns into forest with probability *p*(*f*_*k*_), which depends on the fraction *f*_*k*_ ∈ [0, 1] of neighboring cells occupied by trees. We postulate that this probability peaks at fraction *F*. For simplicity, we assume that *p*(*f*_*k*_) grows linearly from *p*_0_ to *p*_*max*_ as *f*_*k*_ increases from 0 to *F* ; and decreases linearly from *p*_*max*_ to *p*_1_ as *f*_*k*_ increases from *F* to 1:

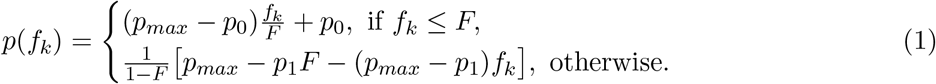

With *F* ≈ 0.5, the probability that an empty cell will regenerate into forest peaks at moderate levels of local forest density, while with *F* close to 1, the above probability peaks at maximum local forest density (i.e., when the neighborhood surrounding a disturbed cell is entirely composed of forest). The latter case assumes that the regeneration of a disturbed cell increases with local forest density, which is analogous to short range seed dispersal.^2^ In contrast, the former case relaxes this assumption, creating a scenario that is analogous to multiscale seed dispersal (Treep *et al*., 2021). We define the amount of new biomass 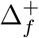 as the number of empty cells regenerated into forest in the current round, and the fraction of cells *f* occupied by trees at the end of the round. Varying parameters (*F, p*_0_, *p*_1_, *p*_*max*_) allows us to examine the effects of different environmental conditions on the model dynamics.

### Agents and their actions

We posit that each agent *i* is characterized by their clearing request *x*_*i*_ ≥ 0 and their sanctioning (acceptability) threshold *y*_*i*_ ≥ 0. The clearing request *x*_*i*_ is the number of cells *i* wants to clear given the actual number of cells occupied by trees. Following the standard convention in the common pool resource literature, we assume that each agent can clear at most *N* -th fraction of the total number of cells occupied by trees (Safarzynska, 2020). However, to clear these cells, *i* needs help from others (Downey *et al*., 2020).

### Dynamics of helping behavior

We assume two factors contribute to the propensity of agent *i* to help *j* with the request *x*_*j*_: normative reasoning and reciprocity. First, *i* tends to reject requests they perceive unacceptable (i.e., those with *x*_*j*_ *> y*_*i*_) and provide help in response to acceptable requests (i.e., those with *x*_*j*_ ≤ *y*_*i*_). Second, helping decisions are also driven by direct reciprocity: agents tend to help more group-mates who have helped them in the past, where the history of help received by *i* from *j* is captured by value *h*_*j,i*_ ∈ [0, 1]. The parameter *µ* ∈ [0, 1] captures the relative importance of normative reasoning in making helping decisions, with respect to the expectation for reciprocity given the helping history of *h*_*j,i*_. As a result, the probability *q*_*i,j*_ that *i* will help *j* (*j≠i*)^3^ with the request *x*_*j*_ is

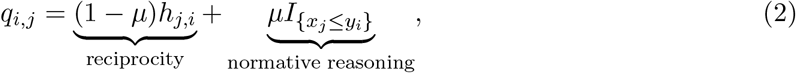

where 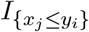 is an indicator function equal to 1 if the clearing request *x*_*j*_ exceeds the threshold *y*_*i*_, and 0 otherwise. Equation 2 implies that with *µ* close to zero, helping behavior is driven primarily by reciprocity. As *µ* increases, helping behavior becomes increasingly conditioned by normative reasoning. Overall, one can treat *y*_*i*_ as personal sanctioning norm, and 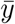 as a social sanctioning norm. Given a fixed relative importance of normative reasoning *µ*, relatively low values of 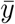 correspond to more severe sanctions, while relatively high values of 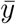 correspond to milder sanctions.

Finally, the history of labor exchange interactions is updated based on the current history and current helping decisions with weights (1 − *α*) and *α* ∈ [0, 1], respectively:

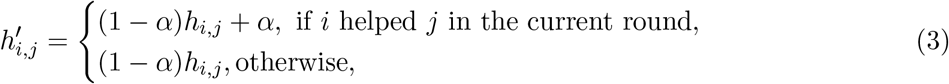

where the prime denotes the next round.

### Individual harvest and payoffs

The individual harvest *x*_*a,i*_ (i.e., the number of cells cleared by agent *i*) is determined by the clearing request *x*_*i*_ and the number of helpers 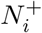. Following Downey *et al*. (2020), we posit that the number of cells *i* is able to clear has a linear relationship with the number of helpers. Specifically, an agent alone (i.e., when 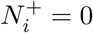) is able to clear *h* cells. Each helper allows an additional *h* cells to be cleared. The individual payoff *π*_*i*_ is determined by the individual harvest *x*_*a,i*_ and total costs *C*_*i*_:

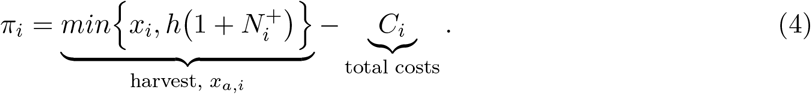

We postulate there are two types of costs in the model. A fixed cost *c*_0_ ≥ 0 is imposed on agent *i≠j* helping another agent *j* and is independent on the number of helpers of *j*. This cost can be treated as the transportation cost or as psychological discomfort from doing something outside of the normal routine. We also assume that each helper of *j* (including *j*) incurs the maximum cost *c ≥* 0 if they clear *h* cells and this cost is proportionally reduced if less than *h* cells are cleared (this depends on the number of helpers 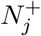). As a result, by helping *j*, agent *i* incurs costs

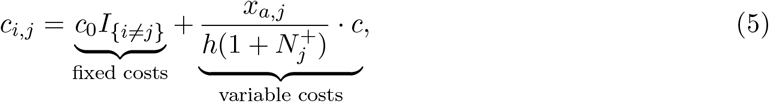

where *I*_*{i≠j}*_ is an indicator function equal to 1 if *i≠j*, and 0 otherwise. Then the total costs *C*_*i*_ are the sum of the costs *c*_*i,j*_ over all agents *j* who received help from *i*.

### Strategy revision protocol

We assume that each round, each agent *i* is independently given an opportunity to revise their clearing request *x*_*i*_ (using myopic optimization) and sanctioning threshold *y*_*i*_ (using payoff-based imitation and innovations) with probabilities *ν*_0_ ∈ [0, 1] and *ν*_1_+*ν*_2_ ∈ [0, 1], respectively. Myopic optimization implies an agent chooses a strategy maximizing their payoff while keeping all others’ decisions the same as in the previous round (Sandholm, 2010; Perry *et al*., 2018; Perry and Gavrilets, 2020). Myopic optimization reflects bounded rationality of agents who are able to maximize their payoff but unable to predict changes in the behavior of others. Here we implement myopic optimization as follows. Agent *i* with the clearing request *x*_*i*_ forms a belief on the number of helpers in the current round based on its expected value given the request *x*_*i*_, the current history of labor exchange interactions, and the previous sanctioning thresholds of others, i.e., 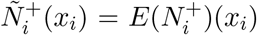. Agent *i* then chooses the clearing request *x*_*i*_ maximizing their material payoff 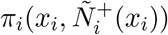 given the expectations on the number of helpers (for more details see Section S1.1 of the SM).

Since individual decisions on sanctioning thresholds affect not the current but next round payoffs and are also conditioned by the choice of clearing requests in the next round, using myopic optimization to revise sanctioning thresholds seems cognitively very difficult – meaning that an agent should make predictions not only about the effects of their current choice on the number of helpers in the next round, but also about their potential clearing request in the next round. To avoid this complexity, we assume agents update their threshold decisions using payoff-based imitation with random innovations (Boyd and Richerson, 1988; Barton, 2014; Creanza *et al*., 2017; Gavrilets and Shrestha, 2021; Lansing, 2009) with probabilities *ν*_1_ and *ν*_2_, respectively (*ν*_1_, *ν*_2_ ≥ 0). To implement errors in the imitation process, we use a Quantal Response Equilibrium-like approach (Goeree *et al*., 2016) with logit errors and a non-negative precision parameter *λ*. Namely, we posit that agent *i* copies the threshold *y*_*j*_ of agent *j* with probability proportional to the j’s payoff *π*_*j*_:

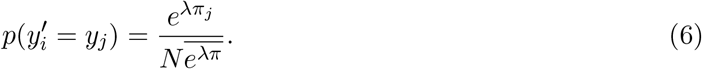

With *λ* → ∞, *i* copies exactly the agent that has the highest payoff; while with *λ* = 0, *i* chooses who to copy completely randomly.

Overall, the effect of the sanctioning (acceptability) threshold *y*_*i*_ on the model dynamics is not trivial. Increasing *y*_*i*_ implies *i* treats more clearing requests as acceptable, which in turn motivates *i* to help others more. On the one hand, this increases costs associated with helping others. On the other hand, it leads to more help received from others in the next round due to reciprocity, but may undermine the ecosystem sustainability. As a result, the interplay of normative reasoning, reciprocity and costs of helping may produce non-trivial effects on the model dynamics. Our goal here is to understand how these factors, along with demographic and environmental parameters, shape the evolution of swidden agriculture. Main model parameters and outcomes are summarized in Table 1. The primary algorithm used in the model is outlined in Figure 1.

**Table 1.** Main parameters and outcomes of the model.

| Parameters | Their meaning |
| --- | --- |
| $\mu$ | importance of normative reasoning |
| $N$ | number of agents |
| $c_0$ and $c$ | cost parameters |
| $F$ | environmental parameter |
| Outcomes | Their meaning |
| $\bar{x}$ | average clearing request |
| $\bar{x}_a$ | average harvest |
| $\bar{y}$ | average sanctioning threshold |
| $\bar{N}^+$ | average number of helpers |
| $\bar{f}$ | remaining fraction of forest |
| $\Delta_f^+$ | new biomass |
| $\bar{\pi}$ | average payoff |

**Figure 1:**
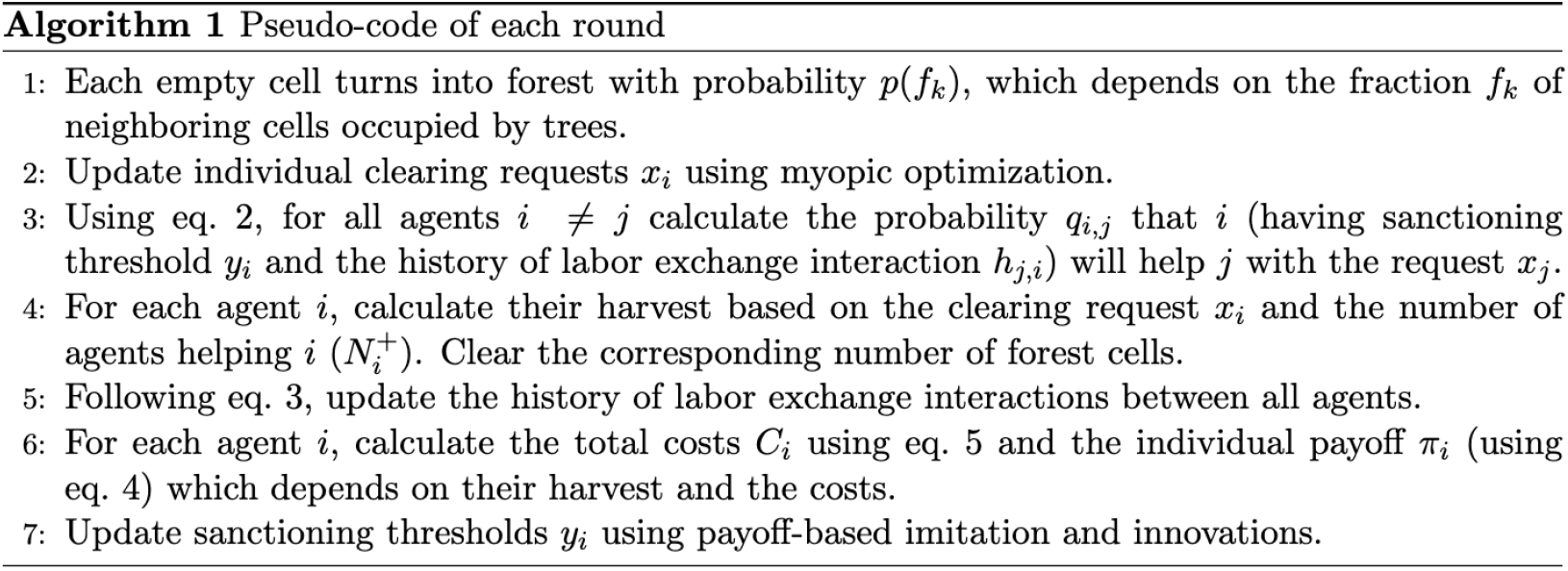
Pseudo-code of each round of the model.

## 3 Results

Some analytical progress is possible for special cases of the model (see the Sections S1.2 - S1.4 of the SM), but to study the overall model dynamics we employ agent-based simulations. We use the following set of default parameter values: *N* = 40, *L* = 120, *T* = 10^3^, *p*_0_ = 0.01, *p*_1_ = 0.06, *p*_*max*_ = 0.12, *F* = 0.5, *h* = 3, *c*_0_ = 0.1, *c* = 0.2, *α* = 0.3, *µ* = 0.75, *λ* = 10, *ν*_0_ = *ν*_1_ = 0.2, and *ν*_2_ = 0.1. Initial clearing requests *x*_*i*_ are drawn randomly and independently from a discrete uniform distribution on *{*1, ‥, 5*}*, while initial sanctioning (acceptability) thresholds *y*_*i*_ are drawn randomly and independently from a uniform distribution on *{*0, ‥, *hN}*. We posit that initially all cells are occupied by tress (i.e., *f*_0_ = 1). All initial helping histories *h*_*i,j*_ are set to zero. These conditions suggest that initially there is no consensus on what size clearing request (*y*) is acceptable, so it can vary from a minimum value of *y*_*i*_ = 0, which indicates agent *i* considers clearing any amount of forest to be completely unacceptable, to a maximum value which is limited by the group size *y*_*i*_ = *hN*, which means agent *i* considers any clearing request to be acceptable. We define outcomes *sustainable* if the average remaining fraction of forest 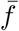 in the last 25% of rounds is greater than 0.05, and *unsustainable* otherwise.

We vary parameters *F, µ, N, c*_0_, and *c* to examine under what conditions sustainable swidden agriculture can evolve. In these settings, three types of swidden regimes are identified.

### 3.1 Three swidden regimes

The low-intensity swidden regime is characterized by very severe sanctions (i.e., 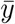 is closed to 0) resulting in no helping behavior (see Figure 2a), so that each agent clears fields individually and a total of approximately *hN* cells are cleared in each round. As a result, the remaining fraction of forest is very high (for the derivation of this equation see Section S1.2 of the SM):

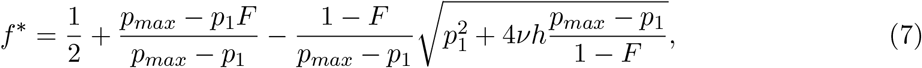

where *ν* is the population density. Clearly, the outcome is characterized by forest conservation, but is inefficient in terms of the agents’ economic productivity, and harvest returns 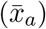 may be insufficient for household needs.

**Figure 2:**
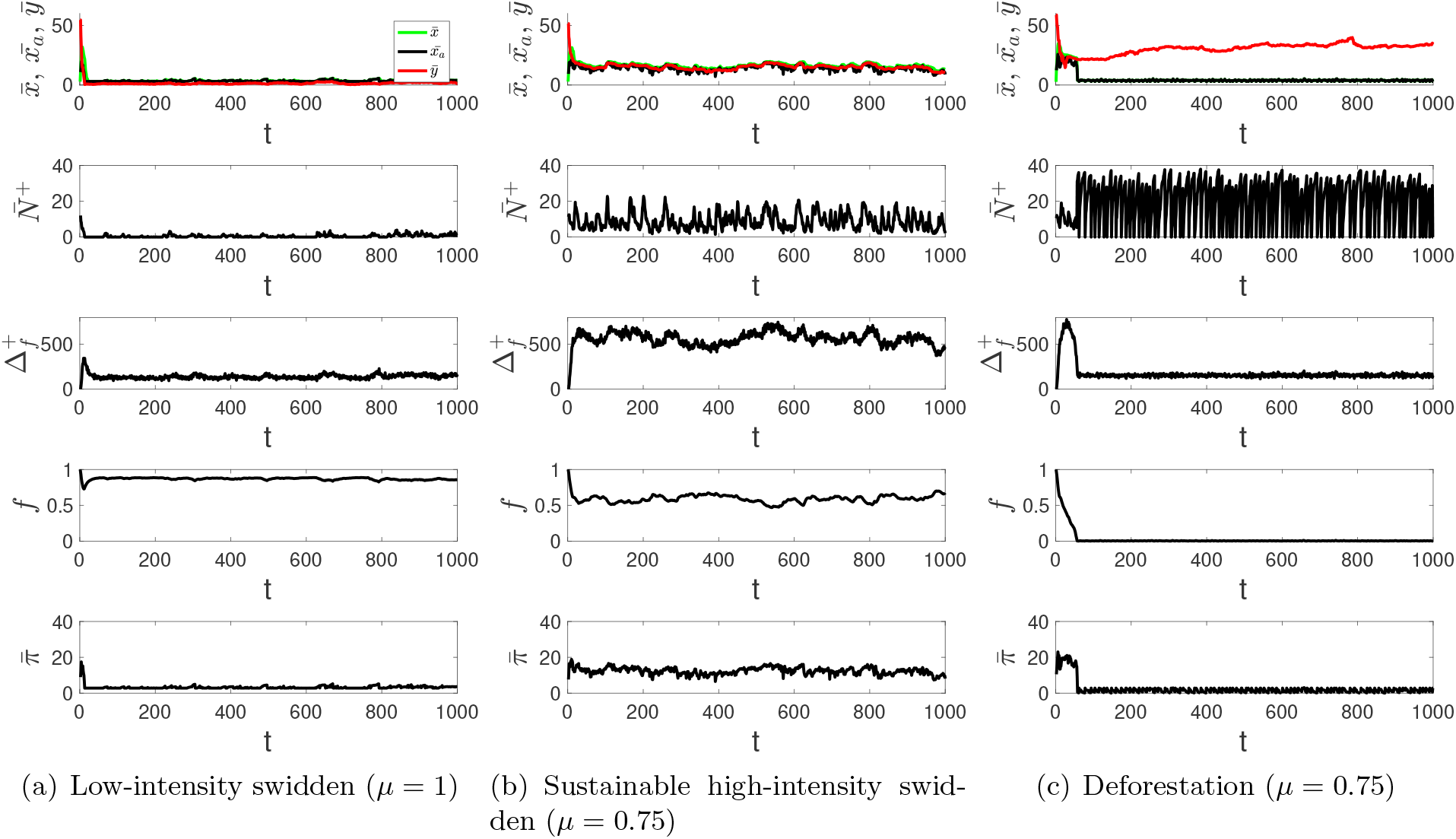
Examples of three swidden regimes. Parameters are at default values, except as indicated.

The sustainable high-intensity swidden regime is characterized by cycles of the sanctioning norm 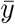, the average harvest 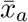, helping behavior 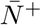, and the remaining fraction of forest *f* (see Figure 2b). Cycling occurs because increasing the norm 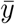 weakens sanctions and promotes helping behavior, which is further reinforced by labor reciprocity when its importance (1 − *µ*) is significantly high. This process is accompanied with a reduction in the remaining fraction of forest, *f*. As the labor exchange network becomes significantly denser, further increases in the threshold *y* are disadvantageous because the associated costs of helping *c* outweigh the potential for increased harvests that results from having more helpers (Figure 3, *t*_1_).^4^ Moreover, it becomes beneficial for agents to increase their clearing requests *x*. This leads to higher harvests due to a dense helping network, and the breakdown of labor exchange connections if the relative importance of normative reasoning *µ* is sufficiently high, which ultimately decreases harvest and limits forest exploitation. In the state with a relatively sparse labor exchange network and relatively severe sanctions (Figure 3, *t*_2_), agents are motivated to reduce their clearing request *x* and/or increase their personal norm *y* to receive more help, however this only works if costs of helping and the relative importance of normative reasoning *µ* are not very high. This tendency leads to an increase in 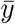 closing the cycle of norms, harvest, helping behavior, and forest dynamics (Figure 3, *t*_3_). Note that the fluctuations in 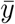 exhibit less variability than the fluctuations in 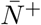 and *x*_*a*_.

**Figure 3:**
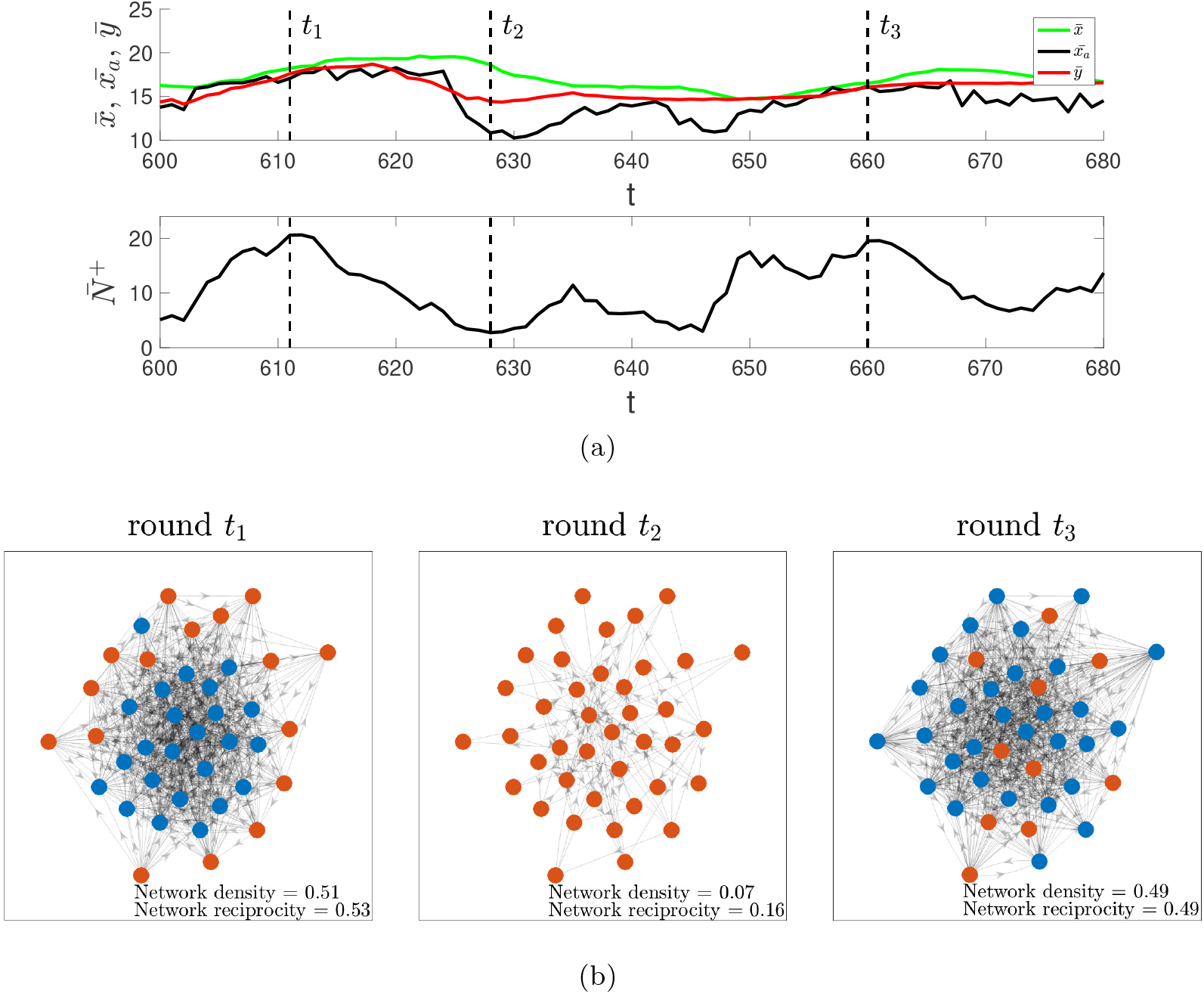
Visualization of cycles of the sanctioning norm, harvest, and helping behavior. (a) Zoomed-in dynamics of the sanctioning norm, harvest, and helping behavior shown in Figure 2b. (b) The corresponding dynamics of the labor exchange network in three rounds: *t*_1_ (relatively loose sanctioning norm 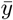 and high compliance with it resulting in a dense labor exchange network), *t*_2_ (relatively tight norm 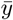 and no compliance with it leading to a sparse labor exchange network driven solely by reciprocity), and *t*_3_ (end of the cycle, similar to *t*_1_). Nodes represent agents and edges correspond to labor exchange interactions between agents. Agents with relatively acceptable clearing requests (i.e., those with 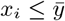) are colored blue, while agents with relatively unacceptable requests (i.e., those with 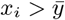) are colored red.

The deforestation regime occurs when *f* ≈ 0 (see Figure 2c). It is characterized by high levels of labor reciprocity and very mild sanctions (i.e., 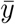 is high) which causes rapid and widespread forest loss.

### 3.2 Effects of social and demographic factors

First, we report the effects of the relative importance of normative reasoning *µ* on the model dynamics and outcomes (Figure 4a). When the relative importance of normative reasoning *µ* is low but non-zero, helping behavior is mainly driven by reciprocity, and the system approaches a unique equilibrium characterized by complete deforestation that is caused by very intense helping behavior. Increasing *µ* leads to bifurcation and bi-stability with the co-existence of two stable equilibria: complete deforestation and sustainable high-intensity swidden agriculture. Around the bifurcation point, sustainable outcomes are characterized by the mildest sanctions and hence the highest harvest and payoffs. As *µ* increases, helping behavior becomes increasingly conditioned by normative reasoning and the frequency of sustainable outcomes increases, but harsher sanctions also evolve, leading to lower harvests and payoffs. As *µ* approaches 1, the sustainable high-intensity swidden regime transforms into a low-intensity swidden regime characterized by no helping behavior, small harvests, strong sanctions, and relatively undisturbed forests. As a result, the sanctioning norm 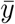 decreases with *µ*, and the remaining fraction of forest *f* exhibits an S-shaped dependence on *µ*^5^. However, the average harvest 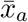, the average number of helpers 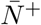, the new biomass 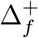, and the average payoff 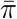 exhibit hump-shaped dependencies on *µ*. This implies the most socially and environmentally efficient outcomes with respect to harvests and biomass are observed at intermediate values of *µ* around the bifurcation point. Overall, the main takeaway is that the balance between normative reasoning and reciprocity is crucial for the evolution of sustainable and intensive swidden agriculture. Indeed, labor exchange is necessary for many of essential tasks associated with swidden agriculture, normative reasoning is necessary to prevent over-harvesting, and reciprocity is necessary to prevent excessive sanctioning.

**Figure 4:**
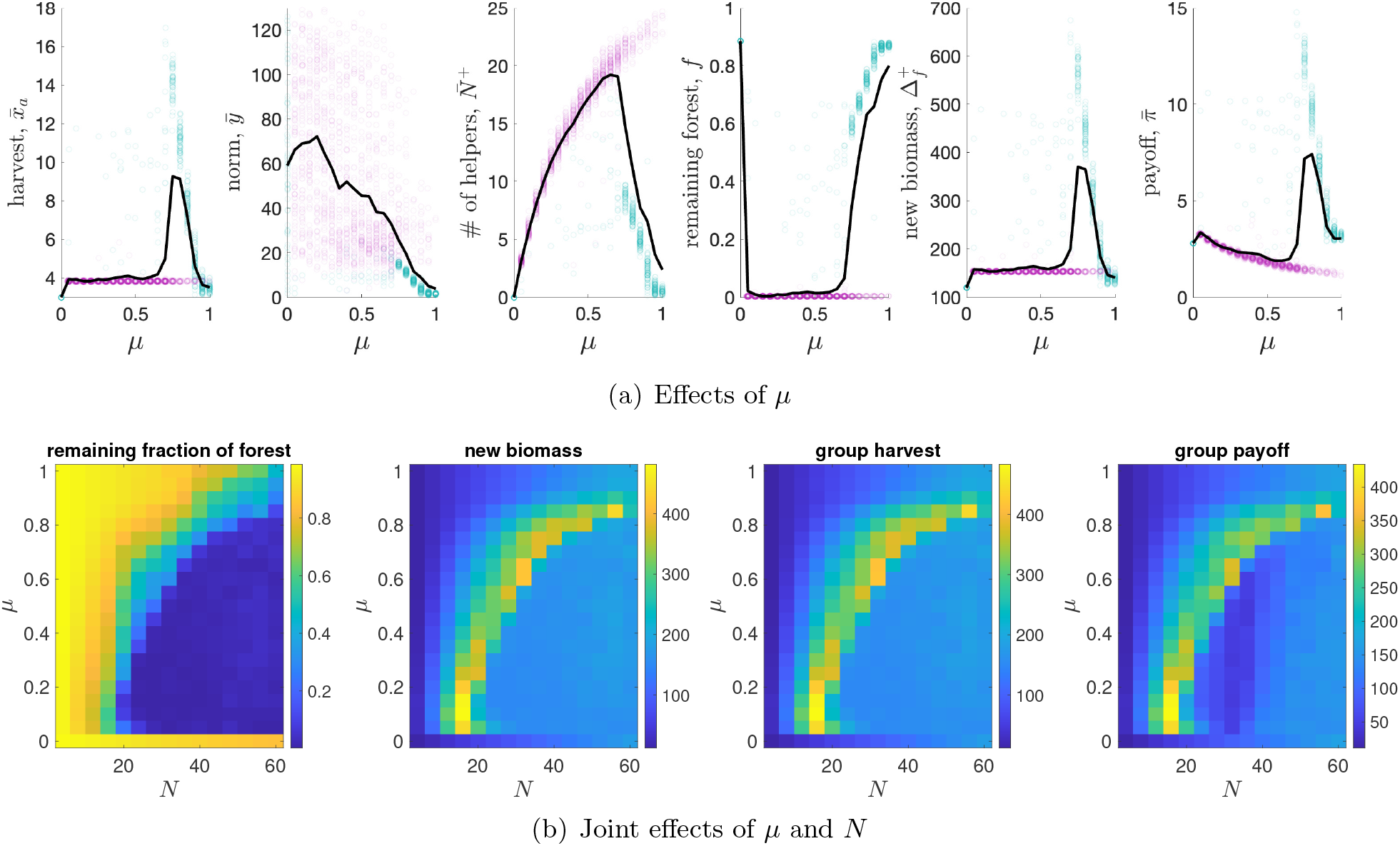
(a) Effects of the relative importance of normative reasoning *µ* on the main model outcomes. Shown are averages in different runs (points) and across all runs (solid black lines). Violet points correspond to unsustainable runs (i.e., those with *f* ≤ 0.05), while green points correspond to sustainable runs (i.e., those with *f >* 0.05). 100 runs each of 1000 rounds for each parameter combination. Results in each run are averages based on the last 250 rounds. All parameters are at default values. (b) Joint effects of *µ* and the group size *N* on: the remaining fraction of forest *f*, the new biomass 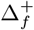, the group harvest *x*_*a*_, and the group payoff *π*. Shown are the averages based on 100 runs each of 1000 rounds for each parameter combination. Results in each run are averages based on the last 250 rounds. Parameters are at default values.

So far, we treated *µ* as a group parameter that applies to all agents in the group. However, within-group behavioral heterogeneity in real groups can significantly affect evolutionary dynamics (Young, 1993; Gavrilets and Richerson, 2017; Neary and Newton, 2017; Efferson *et al*., 2020; Gavrilets, 2020, 2021). To test the robustness of our main result on the key role of the balance between reciprocity and normative reasoning in the evolution of sustainable swidden agriculture, we introduce heterogeneity in the parameter *µ*, so that each individual *µ*_*i*_ is drawn randomly and independently from a normal distribution with mean 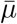 and standard deviation *σ*_*µ*_. As shown in Figure S2 in the SM, a relatively small variation in the parameter *µ* has no effect on the model dynamics. However, a significant variation in *µ* can undermine sustainability and lead to complete deforestation. Overall, a higher importance of normative reasoning is required to maintain sustainable swidden agriculture in a more heterogeneous group.

Finally, we examine the joint effects of group size *N* and the relative importance of normative reasoning *µ* on the model dynamics (Figure 4b). We find that: (1) deforestation is typically observed when *N* is relatively large and *µ* is small; (2) a low-intensity swidden regime evolves if *N* is relatively small and *µ* is large; and (3) a sustainable high-intensity swidden regime is typically observed at the transition between the two other regimes. The results indicate that the sustainable swidden regime is robust to changes in group size and support our previous finding that reciprocity and sanctioning must be balanced in shaping helping behavior to produce sustainable and intensive swidden dynamics, with this balance shifting towards greater importance of normative reasoning as group size increases. Indeed, in small groups, the ability of agents to over-harvest is explicitly limited by group size because the size of the labor exchange network is small. However, increasing *N* creates more opportunities for over-harvesting, so tighter sanctioning norms must evolve to promote sustainable swidden agriculture. Therefore, helping behavior should be driven more by normative reasoning.

### 3.3 Effects of material factors

Here we examine how the cost parameters *c*_0_ and *c* associated with helping behavior shape the evolution of swidden dynamics. The results are shown in Figure S3 in the SM. The main takeaway is that the evolution of sustainable high-intensity swidden agriculture is observed at intermediate values of the total cost parameter (*c* + *c*_0_). Indeed, with relatively small costs of helping, agents are motivated to increase their sanctioning thresholds *y* to expand their labor exchange network and receive higher payoffs, which in turn results in rapid deforestation. In contrast, relatively high costs of helping significantly suppress helping behavior leading to the evolution of severe sanctioning and low-intensity swidden.

We also test the joint effects of the cost parameter *c* and the relative importance of normative reasoning *µ*. As shown in Figure S4 in the SM, the sustainable swidden regime is robust to changes in the structure of material payoffs (e.g., costs), but we predict helping behavior to be more driven by reciprocity as costs increase. The result is intuitive since both costs and the relative importance of normative reasoning are factors controlling the evolution of sanctioning thresholds *y*_*i*_, which should be sufficiently high to promote intensive swidden agriculture and at the same time sufficiently low to maintain sustainability. Therefore increasing costs create material selective pressures suppressing the sanctioning thresholds *y*_*i*_, which should be compensated by social selective pressures promoting an increase in the thresholds that is caused by more reciprocal relations.

### 3.4 Effects of environmental factors

Here we report how the interplay of environmental and social factors shapes the resulting swidden dynamics. First, we test the robustness of our results to changes in the environmental parameter *F*. We compare the model dynamics for *F* = 0.5 and *F* = 0.95, which represent different dispersal assumptions about the level of local forest density at which forest regeneration is most probable for a disturbed patch. As shown in Figures 5a and S12 in the SM and in Section S2.2 of the SM, the main takeaway is that sustainable high-intensity swidden agriculture can evolve in various ecological settings, but is more efficient in terms of harvests (Figure 5a) and new biomass (Figure S12 in the SM) at values of *µ* that lead to an intermediate level of forest disturbance. When *F* = 0.5, an intermediate level of forest disturbance creates better local conditions for forest regeneration than when *F* = 0.95, leading to higher harvests and new biomass. Moreover, for a fixed group size *N* and cost parameters *c* and *c*_0_, a higher importance of normative reasoning *µ* is required to maintain a sustainable swidden regime with increasing parameter *F*. The result is intuitive: agents should be more constrained in their clearing requests in a forest ecosystem exhibiting less ability to recover from severe disturbances, so a higher importance of normative reasoning *µ* is required.

**Figure 5:**
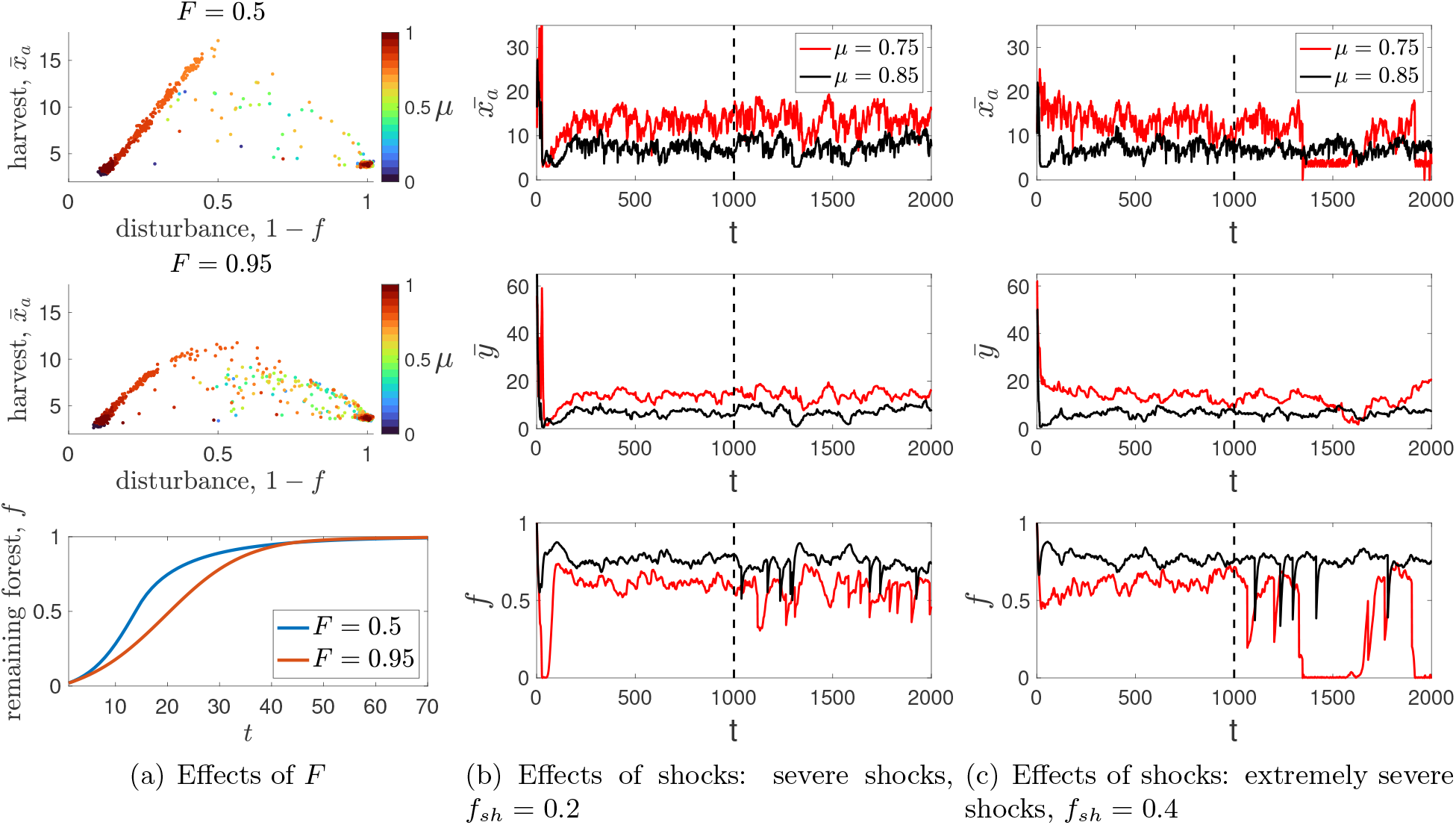
The effects of the environment on swidden dynamics. (a) Top and middle panels: the effects of parameter *F* on equilibrium levels of **harvest** 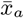 and disturbance (1 − *f*) (the effects on **the amount of new biomass** 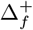 are shown in Figure S12 in the SM). Results are shown for *F* = 0.5 and *F* = 0.95. Each point is the average of 100 runs for each value of *µ* ∈ {0, 0.05, …, 1}. The value of each run is an average of the final 250 rounds out of 1000 total rounds. Other parameters are at default values. Bottom panel: mean-field approximation (see Section S1.2 of the SM) for the reforestation curves in the absence of anthropogenic impact. (b-c) The effects of external shocks on swidden dynamics. Shown are examples of the model dynamics for different values of shock severity *f*_*sh*_ and the relative importance of normative reasoning *µ*. Shocks occur after *t > T/*2 (i.e., to the right to the vertical dashed lines). Parameters: *p*_*sh*_ = 0.01. Other parameters are at default values.

Second, we test the robustness of the sustainable high-intensity swidden regime to external environmental shocks. To model these shocks, we assume that each round, a shock occurs with a small probability *p*_*s*_. This implies *f*_*sh*_*L*^2^ forest cells turn into empty cells, where *f*_*sh*_ ∈ [0, 1] captures the severity of the shocks, and if the remaining number of forest cells is less than *f*_*sh*_*L*^2^, all cells of the grid become empty. Figure 5b,c, Figure S5 in the SM (showing the results for *F* = 0.5), and Figure S11 in the SM (showing the results for *F* = 0.95) indicate that the sustainable high-intensity swidden regime demonstrates considerable resilience to increasing shock severity as long as the shocks are not extremely severe. In harsh environments characterized by extremely severe shocks, a the sustainable high-intensity swidden regime can also be maintained, but the importance of normative reasoning *µ* must increase to ensure the evolution of tighter sanctioning norms in response to shocks.

## 4 Discussion

Many assume that swidden agriculture typically results in deforestation because it is a common property system that is characterized by a lack of centralized regulation (Hardin, 1968). However, the ubiquity and persistence of swidden throughout human history suggests that swidden may be more sustainable than previously understood. Downey *et al*. (2023) proposed that swidden agriculture is best viewed as a complex adaptive system: in the absence of top-down environmental managers or institutions, individual clearing decisions by farmers manifest emergent properties at the landscape scale related to the characteristics and overall state of the forest. Given this proposition, what are the social and ecological mechanisms or factors that could be responsible for maintaining the sustainability of swidden agriculture? The answer lies in the very nature of swidden cultivation: swidden farming involves multiple labor-intensive tasks that cannot be effectively performed individually but only within labor exchange groups, so that farmers depend on receiving help from others. Helping behavior in many small-scale swidden communities tends to be driven by labor reciprocity but also by normative reasoning, that is to say, the process by which farmers’ decide whether to provide help in response to a specific clearing request depending on its perceived acceptability (Downey, 2010; Downey *et al*., 2020; Scaggs and Downey, 2024). For example, in Maya villages in southern Belize, farmers tend to withhold help (i.e., sanction) those who want to clear excessively large fields and tend to provide help to those who want to clear relatively smaller fields. The results of behavioral experiments in these villages demonstrate that such normative reasoning lead to the emergence of a sanctioning norm that might be an important determinant of sustainable swidden agriculture. Here, we attempt to understand how clearing requests, perceptions of acceptability, sanctioning, and forest ecology co-evolve. Our model operationalizes a simple theory for how the social factors that govern labor exchange networks can also determine the emergent sustainable and unsustainable regimes of community forests. In doing this, we shed light on the role of labor reciprocity in this process and how socio-behavioral factors interact with demographic, material, and environmental factors to shape the sustainability of this socioecological system. Finally, we have mapped the full parameter space of possible outcomes related to labor reciprocity, normative reasoning, and several simple ecological models of forest growth, under the assumptions of the model.

Our parsimonious mathematical model of swidden agriculture is built using a minimalist evolutionary approach. Yet, it introduces elements often overlooked in prior modeling research on swidden agriculture, including complex decision-making mechanisms reflecting bounded rationality of agents, and nuanced social dynamics associated with reciprocity and normative reasoning that are derived from ethnographic observations of swidden labor exchange. Three regimes are observed in our simulation results: a low-intensity swidden regime, a deforestation regime, and a sustainable high-intensity swidden regime that maximizes both forest biomass productivity and harvest returns. Deforestation occurs when no significant sanctioning evolves to prevent environmental degradation. In the low-intensity swidden regime, an extremely severe sanctioning norm is observed, which breaks down labor exchange connections leading to small-scale individual swidden farming. As a result, the outcome is environmentally sustainable but it is economically inefficient, and would be unlikely to meet metabolic needs. In the sustainable high-intensity swidden regime, moderate sanctioning evolves, which prevents over-harvesting and encourages labor reciprocity that is essential to swidden agriculture in small-scale societies. This regime is theoretically sustainable for the forest and efficient for society, and we have identified the factors that contribute to the emergence of this sustainable and intensive swidden regime. The key finding is that the balance between normative reasoning and reciprocity in helping behavior is crucial for the evolution of sustainable and intensive swidden agriculture: normative reasoning is needed to prevent over-harvesting, while reciprocity is necessary to prevent excessive sanctioning. Nevertheless, the system does not have to remain in balance over the short term – instead, it is important that agents have the ability adjust their perceptions of allowable land use and clearing requests in order for the system to remain resilient to shocks and to recover from disturbances.

Importantly, our results also indicate that the sustainable high-intensity swidden regime exhibits considerable robustness to changes in demographic, material, and environmental factors. Specifically, it is robust to changes in group size, but agents in larger groups should condition their helping behavior more by normative reasoning and less by reciprocity, so that tighter sanctioning norms can evolve. We also show that excessively small and excessively large groups are both inefficient swidden cultivators. In the former case, agents cannot harvest enough due to lack of sufficient labor partners, and in the latter case forests can become degraded. We find some evidence supporting our results in the ethnographic literature on Q’eqchi’ Maya villages in Belize showing considerable variance in village size with the highest reciprocity rate observed in the smallest village (Downey, 2010). The results also demonstrate considerable flexibility of human behavior to changes in the group size and composition. Indeed, a documented exodus of almost half the population from Graham Greek village in 2008 and subsequent re-population by recruited families from other villages led to increased reciprocity rate, but not to settlement collapse (Downey, 2010).

Finally, we demonstrate that sustainable high-intensity swidden agriculture can evolve in a variety of ecological settings, but that both harvest and biomass productivity are most efficient when the balance between normative reasoning and reciprocity leads to an intermediate level of forest disturbance (Connell, 1978). This is consistent with previous calls highlighting the importance of landscape disturbance for understanding how swidden agriculture is integrated into ecological dynamics (Balée, 2023; Bird, 2015), as well as remote sensing analysis which indicates that tree canopy diversity in Maya community forests is highest at intermediate levels of swidden disturbance (Downey *et al*., 2023). As far as we are aware, ours is the first theoretical study to implement a forest growth model which shows that a combination of reciprocity and normative reasoning can lead to intermediate levels of forest disturbance that enhance both agricultural and ecological productivity. In addition, our results show that the sustainable high-intensity swidden regime exhibits considerable resilience to increasing severity of environmental shocks negatively affecting the forest as long as the shocks are not extremely severe. With extremely severe shocks, a sustainable swidden regime can also be reached, but the likelihood of the settlement collapse is much higher. Alternatively, the relative importance of normative reasoning *µ* needs to be increased to ensure the evolution of tighter sanctioning norms to cope with the extremely severe shocks. This is well in line with ethnographic observations that although forest-damaging hurricanes are quite common in Belize (Friesner, 1993; McCloskey and Keller, 2009), many Maya communities are able to cope with these shocks. Moreover, our theoretical findings support recent work of Stinson and Mcloughlan (2024) suggesting that Maya communities should be able to adapt and recover from the outbreak of climate-change driven wildfires that affected Belize in 2024 (Jones *et al*., 2024) if they are supported in culturally appropriate manner.

In closing, we note that while our model operationalizes complex dynamics related to social norms, swidden labor, and forest ecology in a simple and mathematical way, individual decisionmaking in swidden communities – as indicated by previous ethnographic research – is a much more complex psychological and strategic process. Indeed, determining field sizes, desired harvest levels, and managing one’s labor partnerships in the real world involve considering household composition, economic need, time allocation, limited information, assessment of opportunity costs, and risk. However, we note that it is the purpose of modeling to simplify this realty to better understand the underlying social and ecological dynamics, which we do by focusing on a limited set of factors, labor reciprocity and normative reasoning. Myopic optimization and payoff-based imitation are parsimonious techniques to model processes of strategic behavior, social learning, and adaptation that are significantly more complicated in the real world. Nevertheless, in using these approaches, we illustrate one way it can occur, and in doing so, we hope to better understand the underlying dynamics of swidden agriculture and to provide a plausible demonstration of self-organization in swidden agricultural systems. We also focus attention on modeling social dynamics related to labor reciprocity and normative reasoning, and model forest dynamics in a very minimalist way. This decision was guided by our examination of the swidden modeling literature, which we found to be missing models explaining how social regulation of environmental processes can occur. Modeling a wider range of forest succession stages and the effects of ecological dynamics such as seed dispersal on harvest levels, ecosystem diversity, and productivity would make the model more realistic and potentially useful for understanding real environments (Downey *et al*., 2023), although increasing significantly its complexity. Finally, our current model focuses exclusively on individual decisions about field sizes, however field location and land tenure practices (Barton, 2014) are also important determinants of swidden cultivation. Overall, we believe that incorporating individual decisions on the location of new clearings along with the spatial dependencies of costs, harvest, and forest regeneration will also help explain the real patterns of swidden mosaics, for example as they are observed in remote sensing data (Downey *et al*., 2023). We leave these extensions of the model for future work.

## 5 Conclusion

Swidden agriculture is one of the most widely studied examples of a coupled human and natural system, yet the scarcity of formal models incorporating both its social and environmental dynamics is surprising and consequential. Previous ethnographic fieldwork with Maya communities in Belize, has identified key social dynamics, such as reciprocal swidden labor exchange networks, and ecological dynamics, like forest disturbance that, when combined with concepts from complex adaptive systems theory, capture fundamental aspects of swidden that we believe are relevant to many societies worldwide, in the past, present, and future. In this study, we have distilled a parsimonious model of village-scale social norms related to agricultural labor that interacts with forest disturbance ecology, that is capable of producing a wide range of emergent sustainable and unsustainable outcomes. Our simulation results suggest that the model is resilient across diverse ecosystem types, robust to various perturbations, and that it generates intuitive results like under-utilization and deforestation, as well as counter-intuitive outcomes such as enhanced harvests and biomass productivity (carbon sequestration). These qualitative characteristics are expected in swidden systems, which is a persistent and widespread subsistence strategy throughout human history. We believe our theoretical model may have significant implications for understanding the role that small-scale and Indigenous swidden societies can play in global efforts against climate change. Importantly, our model does not rely on top-down ecosystem management or formal institutions to achieve sustainability. This study focuses on developing and documenting a theoretical model that can be reproduced, tested against empirical observations, and adapted for further scientific research and policy discussions. Overall, we suggest that simple models of swidden agriculture, faithful to observed social and ecological dynamics and able to account for real-world patterns, are essential for advancing our understanding of swidden agriculture and informing policy development in the modern era of climate change.

## Data accessibility

The simulations were performed in MATLAB (Version R2024a). The code for this research work is provided in Section S3 of the SM.

## Research ethics

The model presented in this paper was informed by ethnographic research conducted in Indigenous villages. Permission to conduct the research was obtained from the leaders in both study villages according to customary cultural practices. Ethnographic research permits were obtained from the Belize Institute for Social and Cultural Research (ISCR/H/2/211). The research project was reviewed and approved by the Ohio State University Institutional Review Board (Study Id: 2017B0387).

## Supplementary Materials

### S1 Supplementary analytical results

#### S1.1 Derivation of the best response function

Here we derive the best response function of agents using myopic optimization. Specifically, agent *i* with clearing request *x* forms a belief about the number of helpers in the current round based on its expected value given the request *x*, the current history of labor exchange interactions, and the previous sanctioning thresholds of others, i.e.:

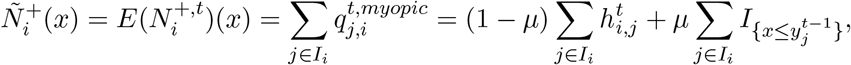

where *I*_*i*_ = *{*1, ‥, *N}\{i}*. Agent *i* then chooses the clearing request *x* maximizing their material payoff 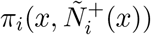 given the expectations on the number of helpers, where

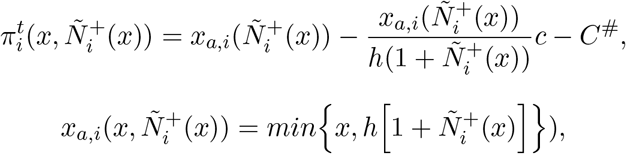

and *C*^#^ are costs that do not depend on *x*. As a result, with *c* = 0 the best response function is

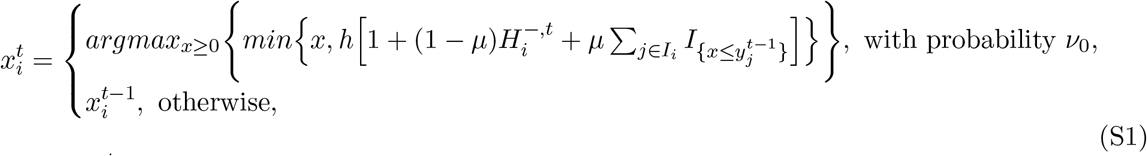

where 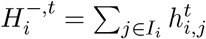 is the history of help provided by agent *i* up to round *t*.

Further, we present supplementary analytical results for some special cases of the model where analytical progress is possible. This allows us to better understand the dynamics of the model and nurture our intuition about the results of agent-based simulations presented in the main text.

#### S1.2 Special case 1: fixed sanctioning norm and no reciprocity

Here we study a special case of the model with no costs (i.e., with *c* = *c*_0_ = 0), a fixed sanctioning norm *Y* ≥ *h* (i.e., assuming all *y*_*i*_ = *Y* = *const*), and helping behavior driven solely by normative reasoning (i.e., with *µ* = 1). This case allows us to deepen our understanding of the forest dynamics, while keeping the social dynamics as simple as possible. For analytical progress, we use a mean-field approximation.

##### Derivation of the mean-field approximation for the forest dynamics

The dynamics of the remaining fraction of forest *f*^*t*^ depends on the fraction of cells turned into forest due to forest regeneration (which we denote as *r*^*t*^) and the fraction of cells cleared by agents (i.e., 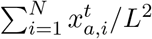):

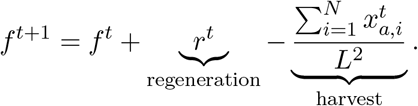

To obtain an explicit form of this equation, we make two simplifying assumptions. First, we assume that the neighborhood topology of each empty cell is similar to the global topology of the grid. This means that the fraction of cells occupied by trees in the local neighborhood of an empty cell in round *t* is *f*^*t*^, implying *r*^*t*^ = *p*(*f*^*t*^) *·* (1 − *f*^*t*^), where *p*(*f*^*t*^) is the probability that an empty cell with fraction *f*^*t*^ forested cells in its local neighborhood will turn into forest. The assumption seems reasonable as agents can choose any cell in the grid, and new cells to clear are chosen randomly among those available.

Second, we make a simplifying assumption about total harvest. Since *Y* ≥ *h*, it is straightforward that agents using myopic optimization to make clearing decisions, choose clearing requests equal to the sanctioning threshold *Y*. As a result:

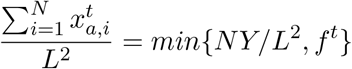

For simplicity, we use the following approximation for the above expression:

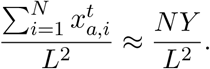

Then, the mean-field approximation for the equation governing the dynamics of forest *f*^*t*^ is:

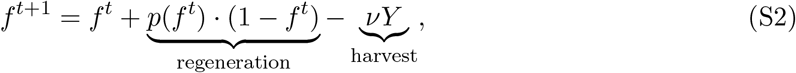

where *ν* = *N/L*^2^ is the population density. Then an equilibrium *f* ^∗^ is a solution to:

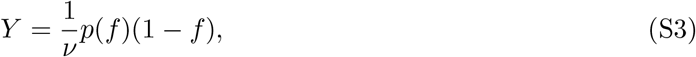

which is Lyapunov-stable if

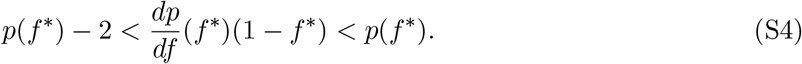

We say that an equilibrium *f* ^∗^ is sustainable if *f* ^∗^ *>* 0 and unsustainable if *f* ^∗^ = 0. With the piece-wise linear functional form of *p*(*f*) defined by Equation 1, the conditions for the existence and stability of a sustainable equilibrium are formulated as follows.

###### Proposition 1

*A sustainable equilibrium of the system S2, given the piece-wise linear functional form of p*(*f*) *defined by Equation 1*,

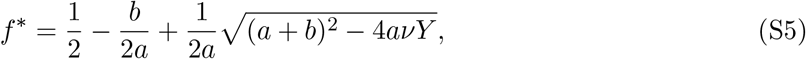

*exists and is stable in three cases*.

(1) 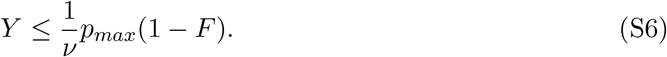

*In this case*, 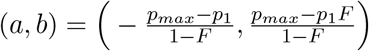 *and f* ^∗^ ∈ [*F*, 1].

(2) 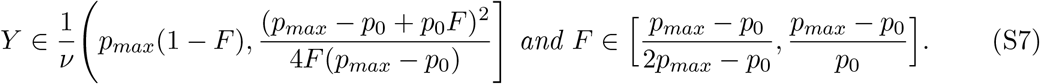

(3) 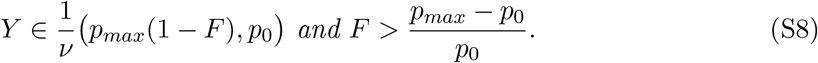

*In the latter two cases*, 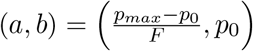 *and f* ^∗^ ∈ [0, *F*).

*Otherwise, the only outcome observed in the model is deforestation*.

The proof of this proposition can be found in section S1.4.

###### Corollary 1

*Let p*_*max*_ *>* 2*p*_0_. *The unique sustainable equilibrium of the system S2 with piece-wise linear functional form of p*(*f*) *defined by Equation 1 characterized by the remaining fraction of forest f* ^∗^ ≥ *f*_*min*_ *exists and is stable if Y* ≤ *Y*_*max*_, *where*

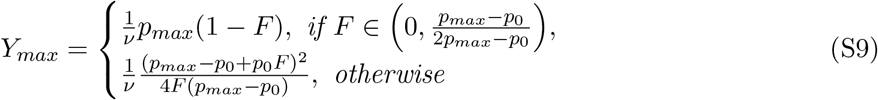

*And*

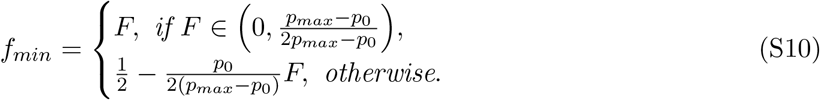

The main takeaway is that there is a maximum sanctioning threshold *Y*_*max*_ beyond which the sustainable equilibrium disappears and only deforestation is observed. Therefore, *Y*_*max*_ can be viewed as the socially optimum outcome when sanctioning is as loose as possible and harvest is as high as possible to ensure sustainability. This results in the remaining fraction of forest *f*_*min*_, which is the minimum fraction of forest that can be observed in a sustainable regime. In addition, it is straightforward to show that *Y*_*max*_ decreases with *F*. The result is intuitive: a stronger sanctioning norm is required in a forest ecosystem exhibiting less ability to recover from relatively more severe disturbances to ensure a sustainable outcome. So naturally, better outcomes for agents are observed when *F* is smaller.

Interestingly, the lowest sustainable fraction of forest *f*_*min*_ exhibits a hump-shaped dependence on *F* peaking at the value of

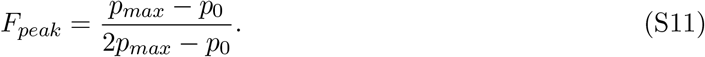

Moreover, with *p*_*max*_ *>> p*_0_, *F*_*peak*_ ≈ 0.5 and for all *F* ∈ [*F*_*peak*_, 1) : *f*_*min*_ ≈ 0.5. This means that if forest regeneration in deforested areas is sufficiently difficult, at least about half of the cells should be occupied by tress in a sustainable equilibrium.

#### S1.3 Special case 2: fixed sanctioning norm, both reciprocity and normative reasoning contribute to helping behavior

Here we study a special case of the model with no costs (i.e., with *c* = *c*_0_ = 0) and a fixed sanctioning norm *Y >* 0 (i.e., assuming all *y*_*i*_ = *Y* = *const*). This case allows us to deepen our understanding of the cycles of helping behavior observed in agent-based simulations.

##### Derivation of the equations governing the model dynamics

For simplicity, we neglect the fact that the clearing request *x* must be an integer and assume that agent *i* updates their decisions in round *t*. Then, following Equation S1, we observe

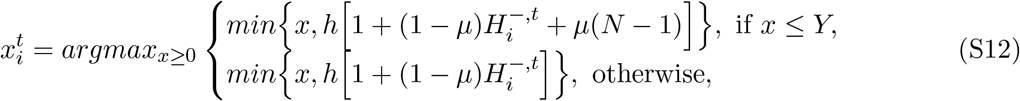

which entails

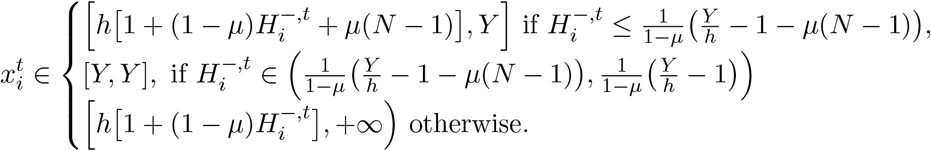

Without loss of generality, the above expression can be reduced to:

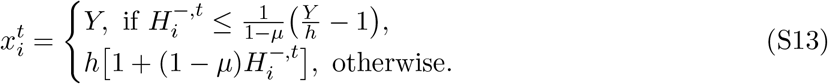

Given Equation 3, the dynamics of the history of labor exchange interactions between agents *i* and *j* can be approximated as follows:

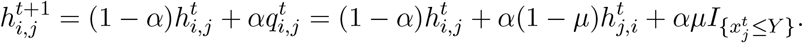

Note that these *N* ^2^ equations can be reduced to 2*N* equations:

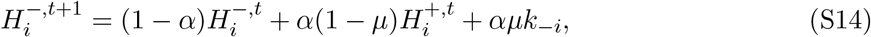

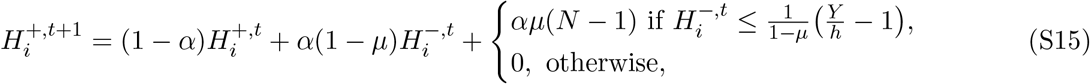

where 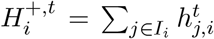 is the history of help received by agent *i* up to round *t* and *k*_−*i*_ is the number of agents in the set *I*_*i*_ with clearing requests not exceeding the threshold *Y*. Since *Y* is the same for all agents in the group, it can be treated as a group sanctioning norm. Therefore, agents with *x* ≤ *Y* can be treated as rule-followers, while those with *x > Y* can be viewed as rule-breakers. We will use this terminology throughout this section.

Overall, the dynamics of the system can be approximated by a system of (3*N* + 1) difference equations: *N* Equations S13 for the clearing requests dynamics, 2*N* Equations S14-S15 for the dynamics of the labor exchange network, and one equation for forest dynamics (see the previous section for the derivation of the mean-field approximation):

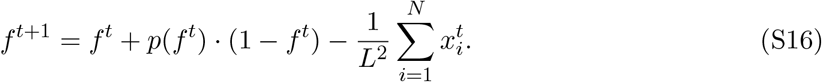

##### Equilibrium regimes

To describe all sustainable equilibria existing in the model, we formulate the following proposition. Let *D* be the set of all *z* ∈ *R*:

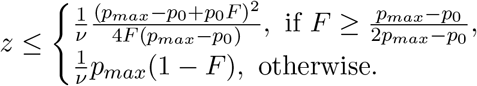

###### Proposition 2

*Let p*_*max*_ *>* 2*p*_0_. *Then the following sustainable equilibria exist in the model:*

(1) *An equilibrium with all rule-breakers exists if*

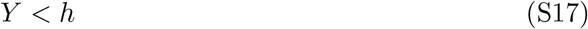

*and*

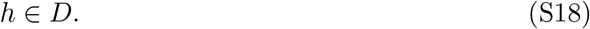

*In this equilibrium, each agent makes a clearing request* 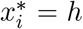 *and no exchange of labor is observed*.

(2) *An equilibrium with all rule-followers exists if*

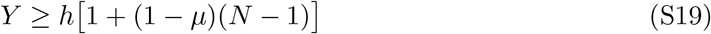

*and*

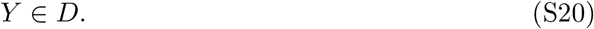

*In this equilibrium, each agent makes a clearing request* 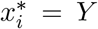, *and the labor exchange network is a complete graph on N nodes*.

(3) *An equilibrium with k* ∈ *{*1, ‥, *N* − 1*} rule-followers exists if*

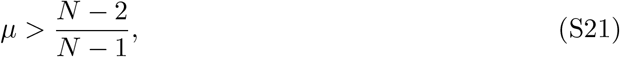

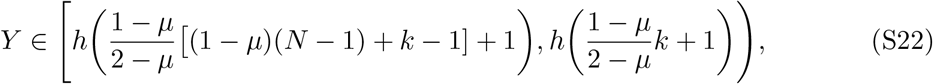

*and*

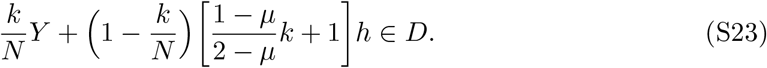

*In this equilibrium, each rule-follower makes a clearing request* 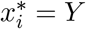, *while each rule-breaker makes a clearing request* 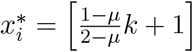.

The proof of this proposition can be found in section S1.4.

First, note that equilibria with *k* ∈ *{*1, ‥, *N* − 1*}* rule-followers and (*N* − *k*) rule-breakers exist in a very narrow parameter region with the relative importance of normative reasoning *µ* very close to 1 and the sanctioning norm *Y* ∈ [*Y*_*low*_, *Y*_*high*_], where 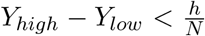. As a result, we exclude these equilibria from further analysis and focus on two sustainable equilibria: with all rule-followers and all rule-breakers, respectively. As shown in Figure S1, the equilibrium with all rule-breakers (low-intensity swidden regime) exists if sanctioning is very severe (i.e., *Y* is low) so that agents have no incentive to help others. The equilibrium with all rule-followers exists in a fairly narrow region characterized by intermediate sanctioning and high importance of normative reasoning (i.e., when *µ* is close to 1). For all other parameter values either no sustainable equilibria exist or cycles of helping behavior are observed. These approximation results for a special case of the model give some insight into why non-equilibrium dynamics are so widespread in our model.

**Figure S1:**
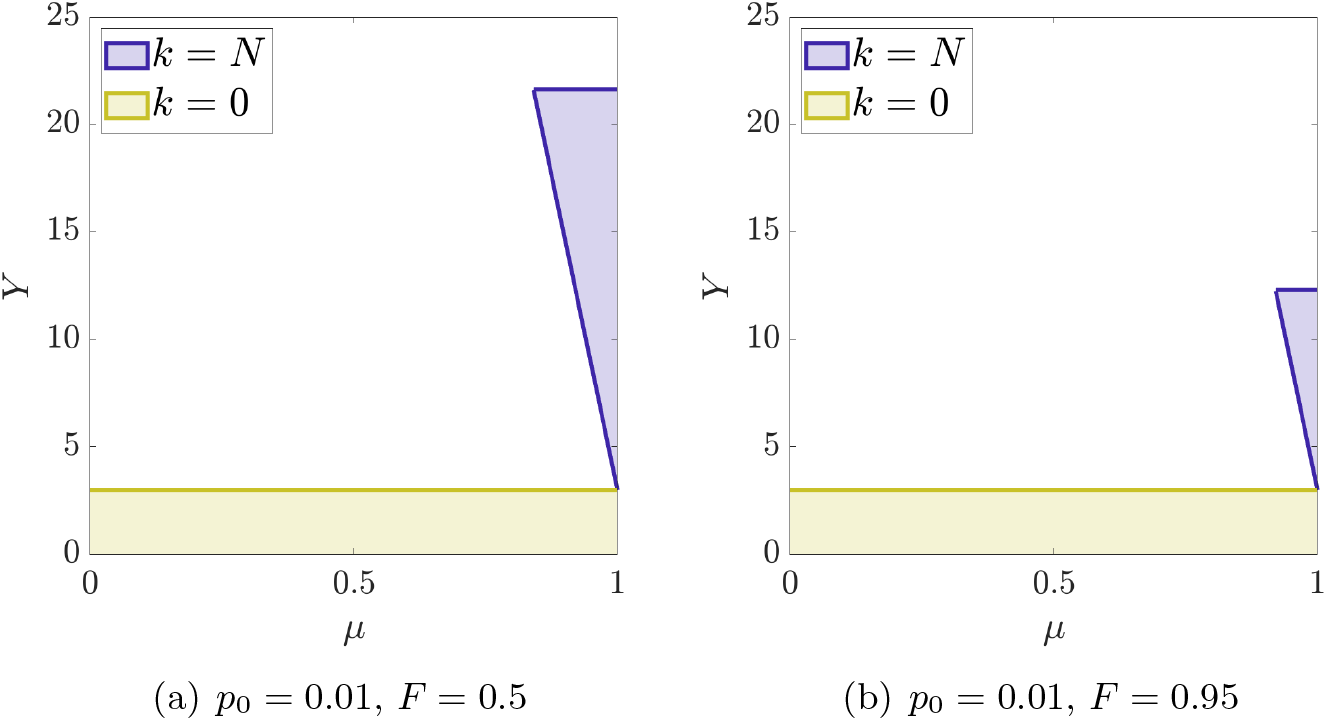
Regions of existence of sustainable equilibria with *k* rule-followers and (*N* − *k*) rule-breakers as a function of the relative importance of normative reasoning *µ* and the sanctioning norm *Y*. Other parameters: *N* = 40, *L* = 120, *h* = 3, and *p*_*max*_ = 0.12.

### S1.4 Proofs

Here we present proofs of all propositions formulated above.

**Proof of Proposition 1**. Let *f* ^*\**^ be a sustainable equilibrium. Note that if *f* ^*\**^ *< F* or *f* ^*\**^ *> F, p*(*f*) is linear in a local neighborhood of *f* ^*\**^, i.e., it has the form *p*(*f*) = *af* + *b*, where

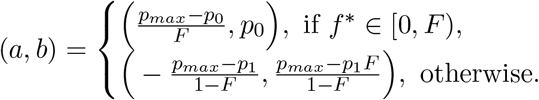

The equilibrium remaining fraction of forest *f* ^*\**^ is a solution to Equation S3, which is quadratic:

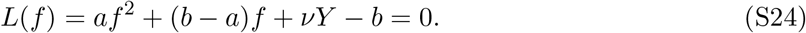

According to Condition S4, *f* ^*\**^ is Lyapunov-stable if

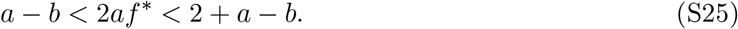

Further, we consider two cases with *f* ^*\**^ *< F* and *f* ^*\**^ *> F* separately.

*The case of f* ^*\**^ *> F*. Since 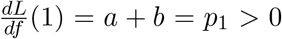 and 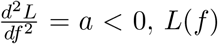, *L*(*f*) is a monotonically increasing function on [0, 1]. Moreover, since *L*(1) = *νY >* 0 and by the Intermediate value theorem, *L*(*f*) = 0 on [*F*, 1] if and only if *L*(*F*) ≤ 0. Moreover, the solution *f* ^*\**^ to *L*(*f*) = 0 on [*F*, 1] is unique and has the form S5. The condition for the existence of this equilibrium, *L*(*F*) ≤ 0, entails Condition S6.

Following Condition S25, it is straightforward to check that *f* ^*\**^ is Lyapunov-stable if

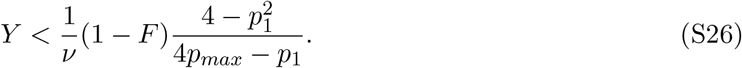

However, Conditions S6 and S26 reduce to Condition S6, so statement (1) of Proposition 1 is proved.

*The case of f* ^*\**^ *< F*. Following Equation S24, one concludes that there can be two steady states of the system S2: 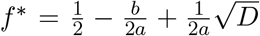 and 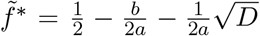, where *D* = (*a* + *b*)^2^ − 4*aνY*. According to Condition S25, the former is stable if 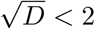, while the latter is unstable. Moreover, *f* ^*\**^ exists if *f* ^*\**^ ∈ (0, *F*) and *D* ≥ 0. Combining these conditions with the condition for local stability, one obtains:

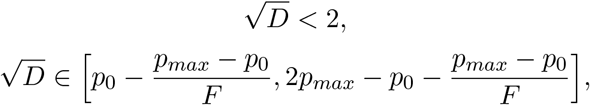

and

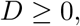

which can be reduced to the last two conditions, which in turn, after straightforward algebraic manipulations, yield statements (2) and (3) of Proposition 1. As a result, Proposition 1 is proved.

**Proof of Proposition 2**. (1). Equations S13 imply all agents are rule-breakers if

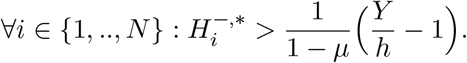

As a result, Equations S14-S15 governing the dynamics of the labor exchange network can be reduced to

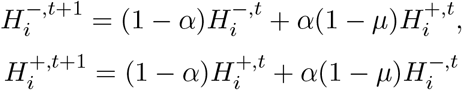

with the steady state 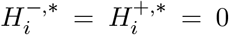 (so there are no labor exchange interactions between agents). Inserting this into Equations S13, we conclude that 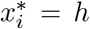 and *Y < h* to ensure the existence of the equilibrium with all rule-breakers. Condition S18 for sustainability stems from Proposition 1. (2) Equations S13 imply all agents are rule-followers if

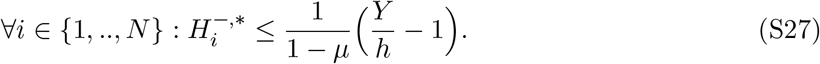

In this case,

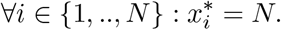

Given this, Equations S14-S15 governing the dynamics of the labor exchange network can be reduced to

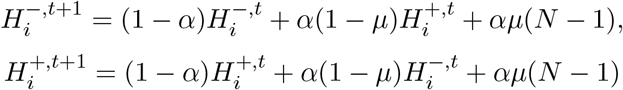

with the steady state 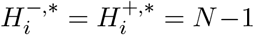 (so that each agent helps all the group-mates). Inserting this into Inequality S27, we obtain Condition S19. Then, Condition S20 for sustainability stems from Proposition 1. (3). Equations S13 imply for each rule-follower

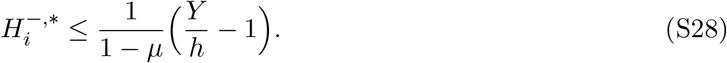

Given this, Equations S14-S15 governing the dynamics of the labor exchange network can be reduced to

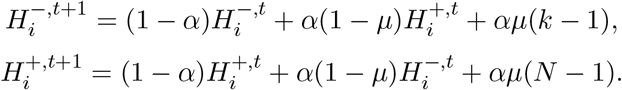

Then, in the steady state

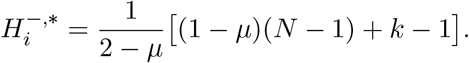

Inserting this expression into Inequality S28, we obtain the lower boundary in Condition S22.

Similarly, Equations S13 imply for each rule-breaker

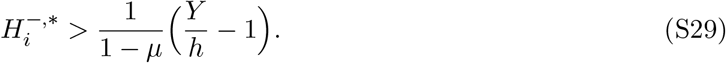

Given this, Equations S14-S15 governing the dynamics of the labor exchange network can be reduced to

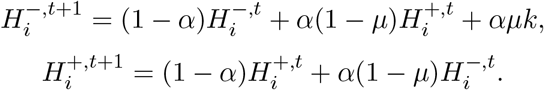

As a result, in the steady state

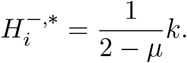

Plugging this expression into Equations S13, we obtain that for each defector, 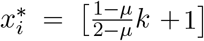. Moreover, plugging this expression into Inequality S29, we obtain the upper boundary in Condition S22. The lower boundary is lower than the upper boundary if

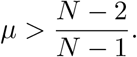

Finally, since the total harvest is 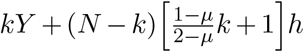, and following Proposition 1, we obtain Condition S23 for sustainability. The proposition is proved.

## S2 Supplementary Figures

**Figure S2:**
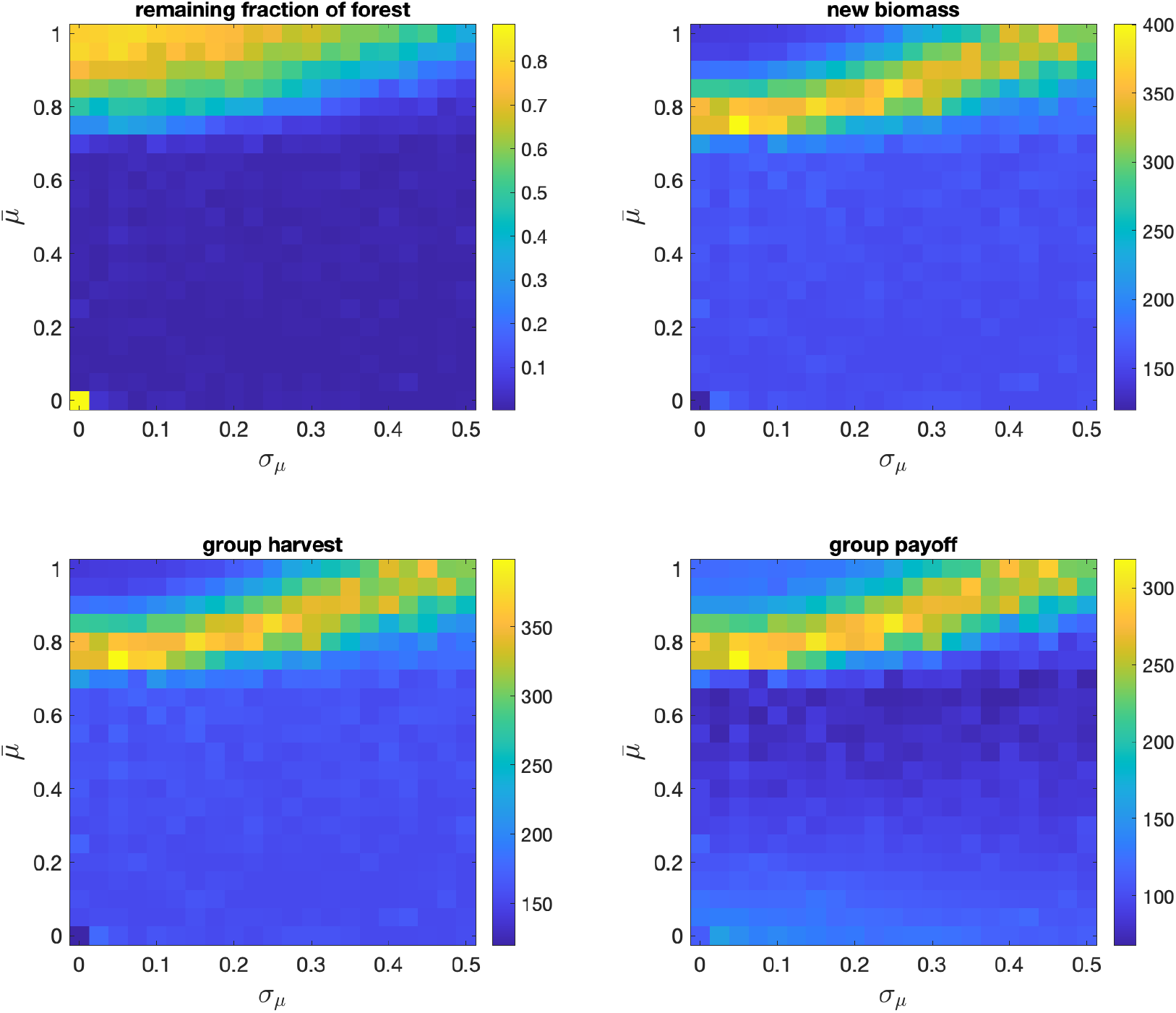
Model outcomes under the assumption of a normal distribution of the relative importance of normative reasoning *µ* with mean 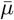 and standard deviation *σ*. Shown are the averages based on 100 runs each of 1000 rounds for each parameter combination. Results in each run are averages based on the last 250 rounds. Parameters are at default values.

**Figure S3:**
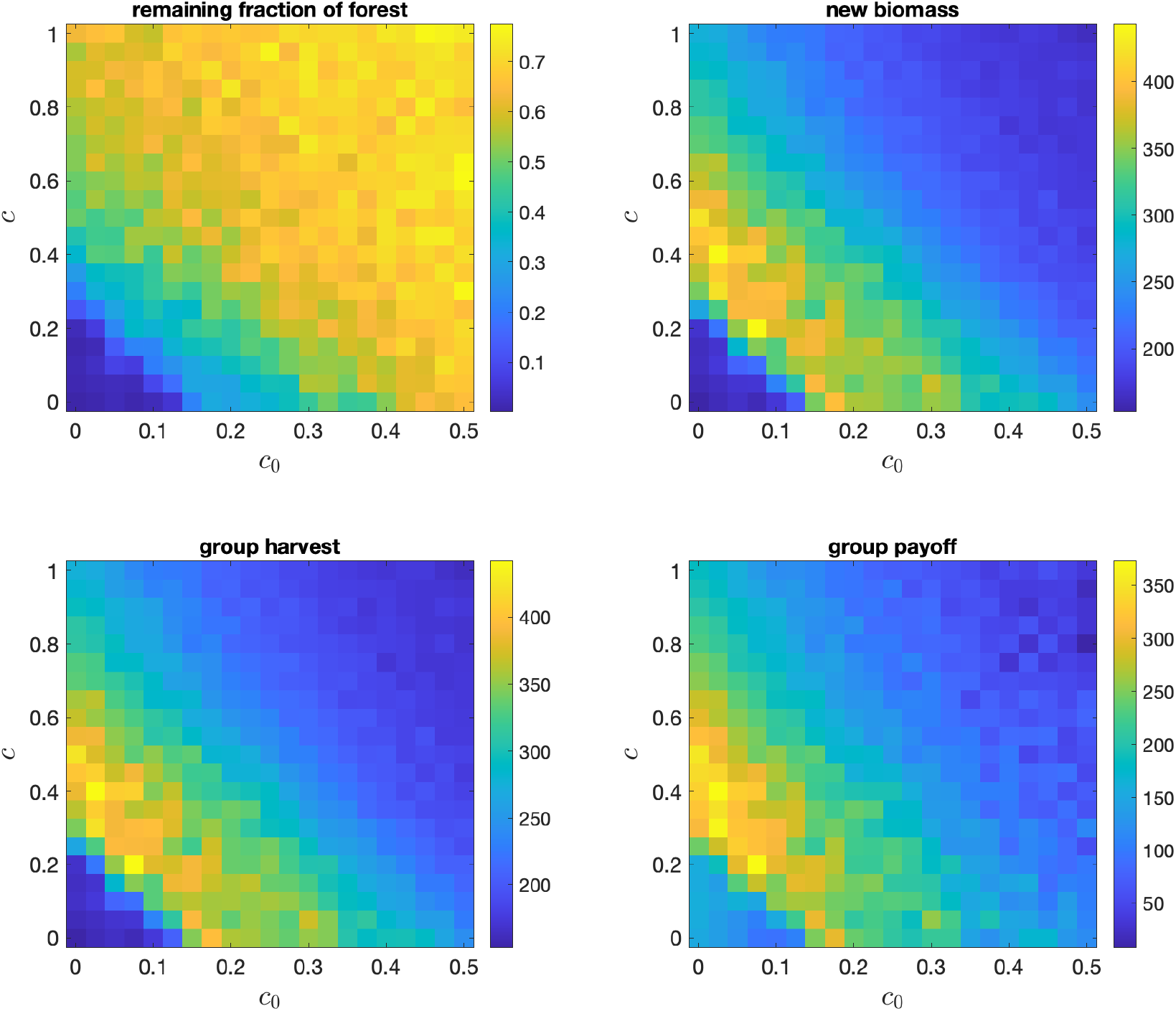
Effects of the cost parameters *c*_0_ and *c* on: the remaining fraction of forest *f*, the new biomass 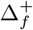, the group harvest *x*_*a*_, and the group payoff *π*. Shown are the averages based on 100 runs each of 1000 rounds for each parameter combination. Results in each run are averages based on the last 250 rounds. Parameters are at default values.

**Figure S4:**
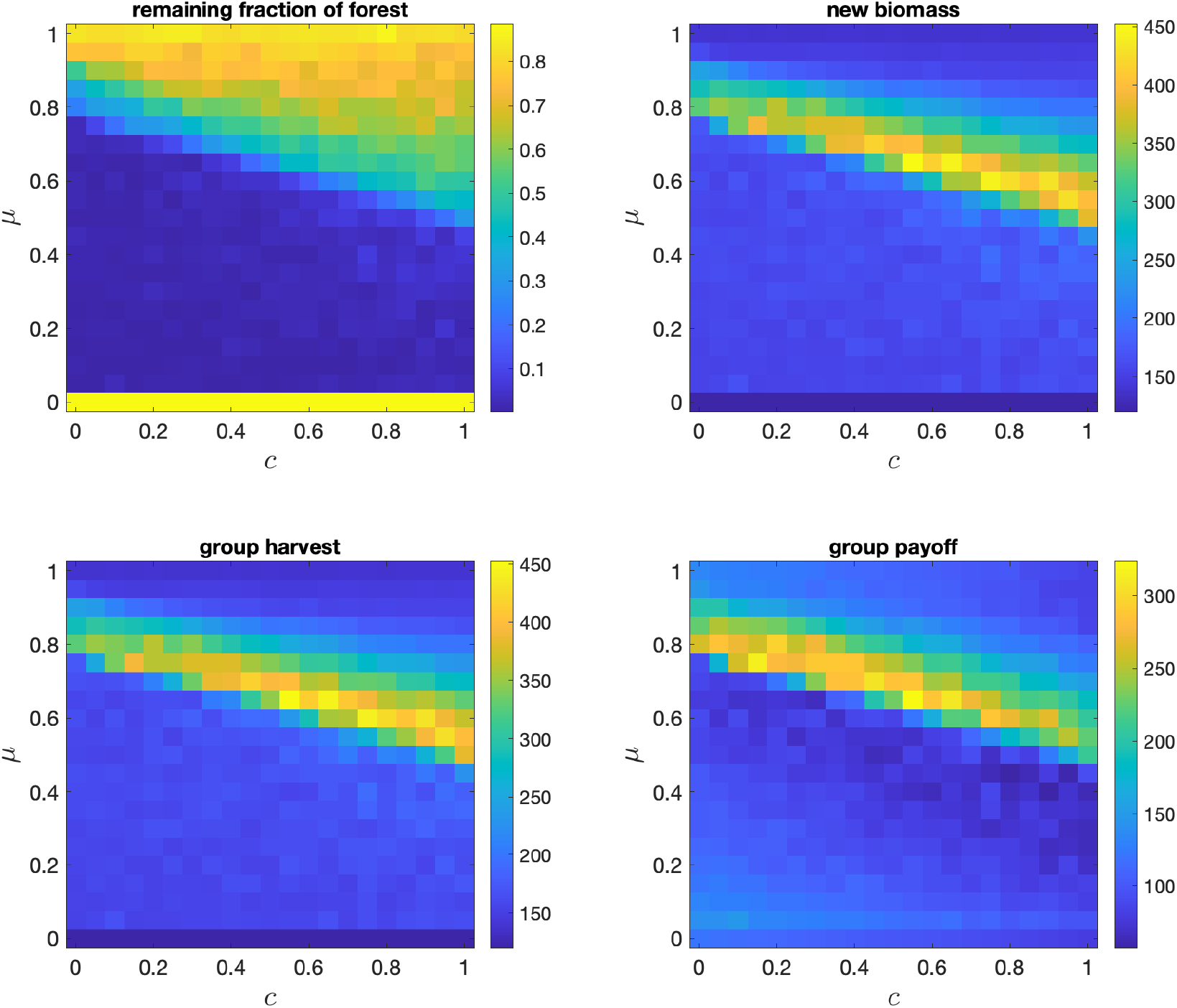
The joint effects of the cost parameter *c* and the relative importance of normative reasoning *µ* on: the remaining fraction of forest *f*, the new biomass 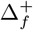, the group harvest *x*_*a*_, and the group payoff *π*. Shown are the averages based on 100 runs each of 1000 rounds for each parameter combination. Results in each run are averages based on the last 250 rounds. Parameters are at default values.

**Figure S5:**
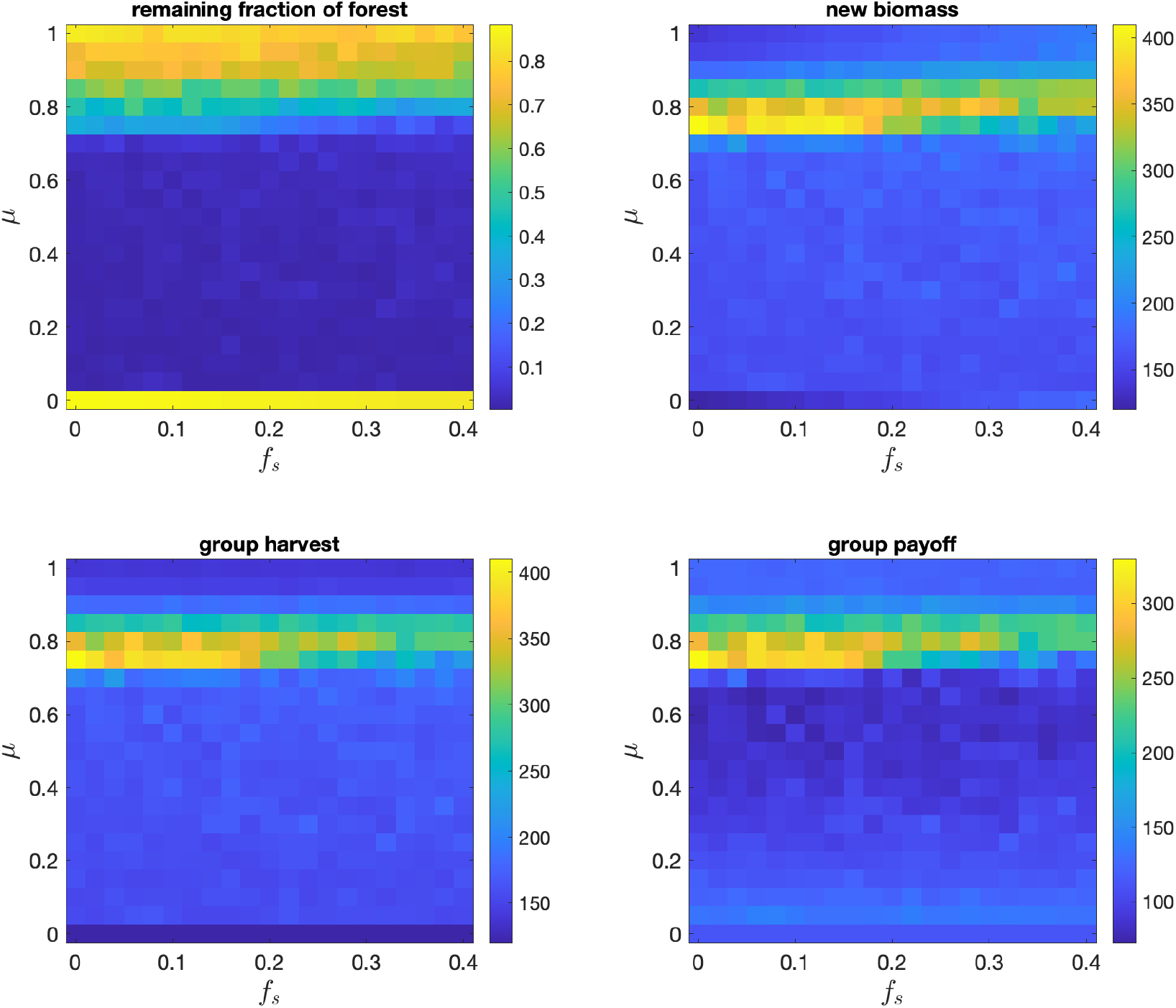
The effects of the relative importance of normative reasoning *µ* and the severity of shocks *f*_*s*_ on: the remaining fraction of forest *f*, the new biomass 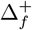, the group harvest *x*_*a*_, and the group payoff *π*. Shown are the averages based on 100 runs each of 1000 rounds for each parameter combination. Results in each run are averages based on the last 250 rounds. Each round, a shock occurs randomly and independently with probability *p*_*sh*_ = 0.01. Other parameters are at default values.

### S2.1 Supplementary Figures for the case of *F* = 0.5

**Figure S6:**
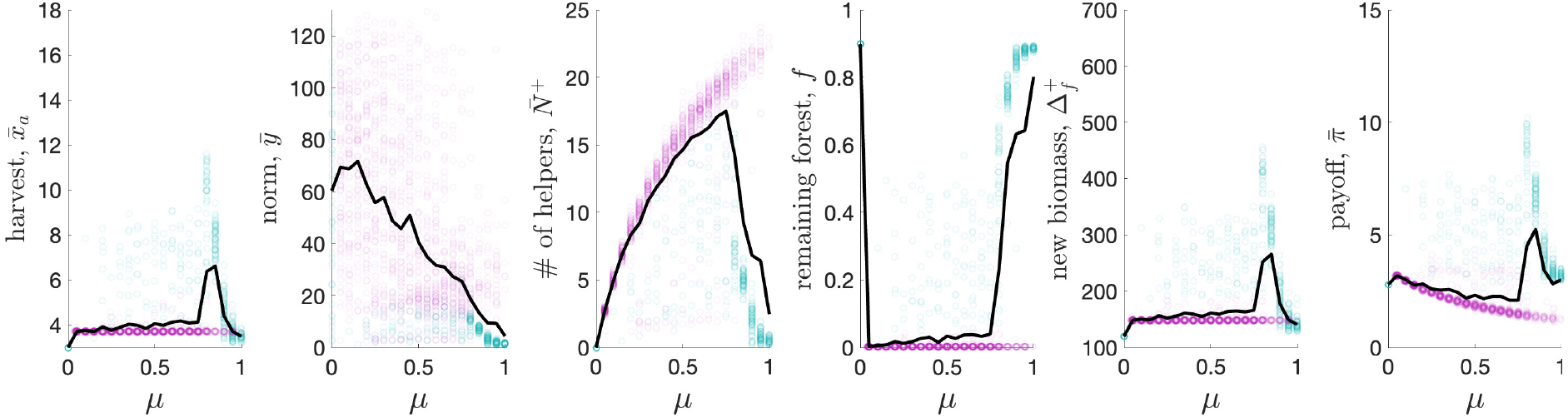
Effects of the relative importance of normative reasoning *µ* on the main model outcomes. Shown are averages in different runs (points) and across all runs (solid black line). Violet points correspond to unsustainable runs (i.e., those with *f* ≤ 0.05), while green points correspond to sustainable runs (i.e., those with *f >* 0.05). 100 runs each of 1000 rounds for each parameter combination. Results in each run are averages based on the last 250 rounds. *F* = 0.95, other parameters are at default values. **The results for** *F* = 0.5 **are shown in Figure 4a**.

**Figure S7:**
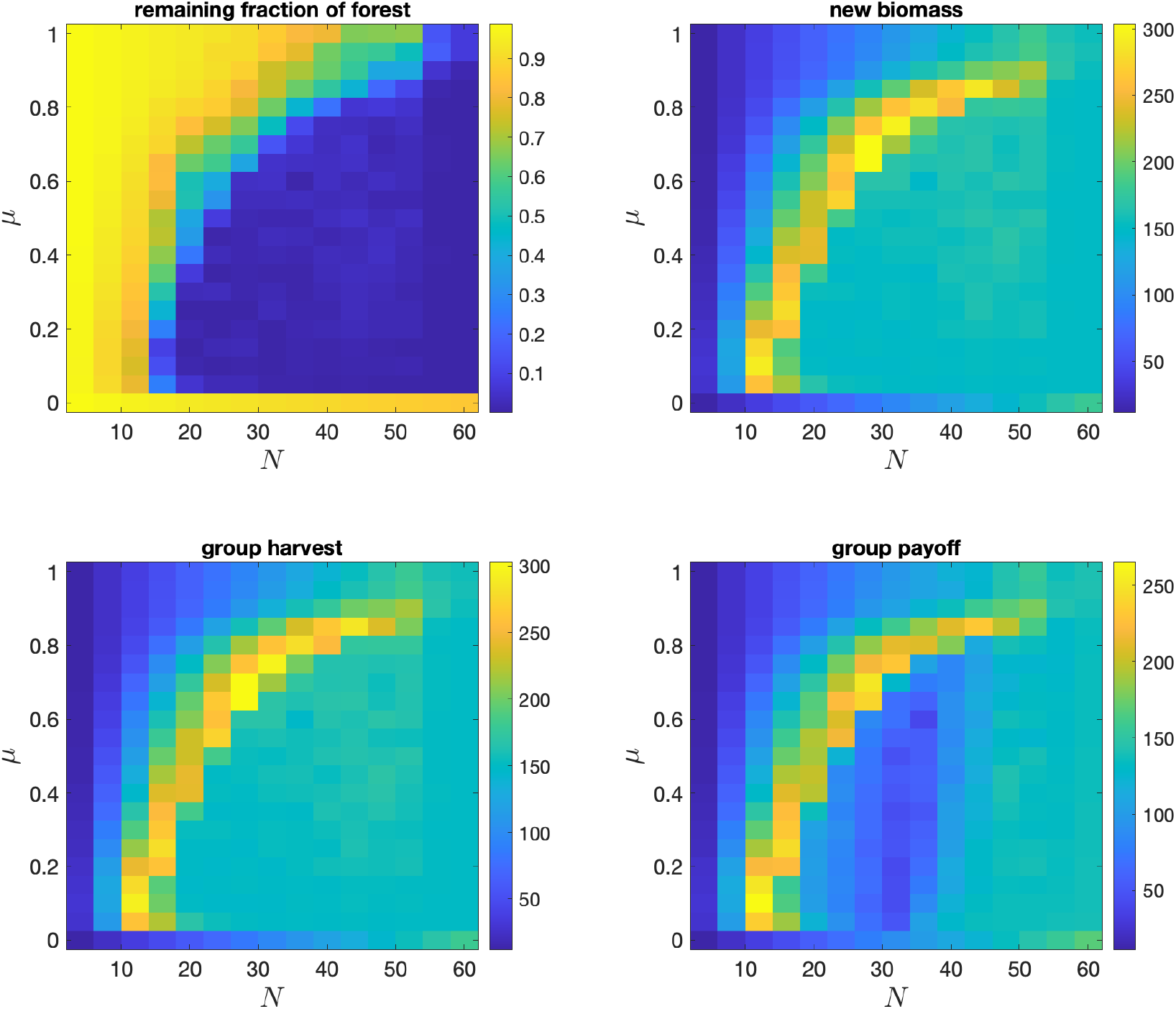
Joint effects of the relative importance of normative reasoning *µ* and the group size *N* on: the remaining fraction of forest *f*, the new biomass 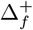, the group harvest *x*_*a*_, and the group payoff *π*. Shown are the averages based on 100 runs each of 1000 rounds for each parameter combination. Results in each run are averages based on the last 250 rounds. *F* = 0.95, other parameters are at default values. **The results for** *F* = 0.5 **are shown in Figure 4b**.

**Figure S8:**
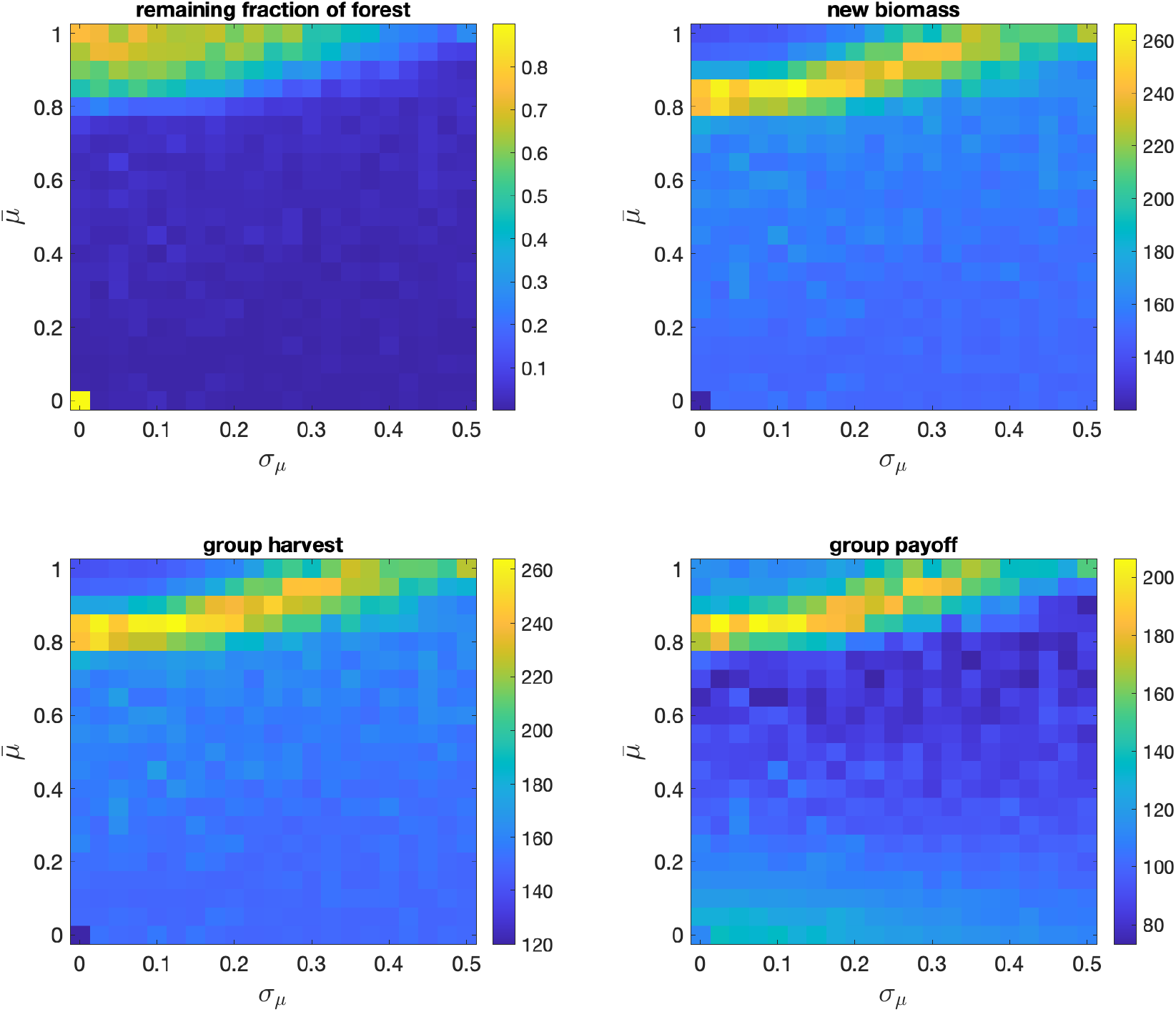
Model outcomes under the assumption of a normal distribution of the relative importance of normative reasoning *µ* with mean 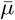 and standard deviation *σ*. Shown are the averages based on 100 runs each of 1000 rounds for each parameter combination. Results in each run are averages based on the last 250 rounds. *F* = 0.95, other parameters are at default values. **The results for** *F* = 0.5 **are shown in Figure S2**.

**Figure S9:**
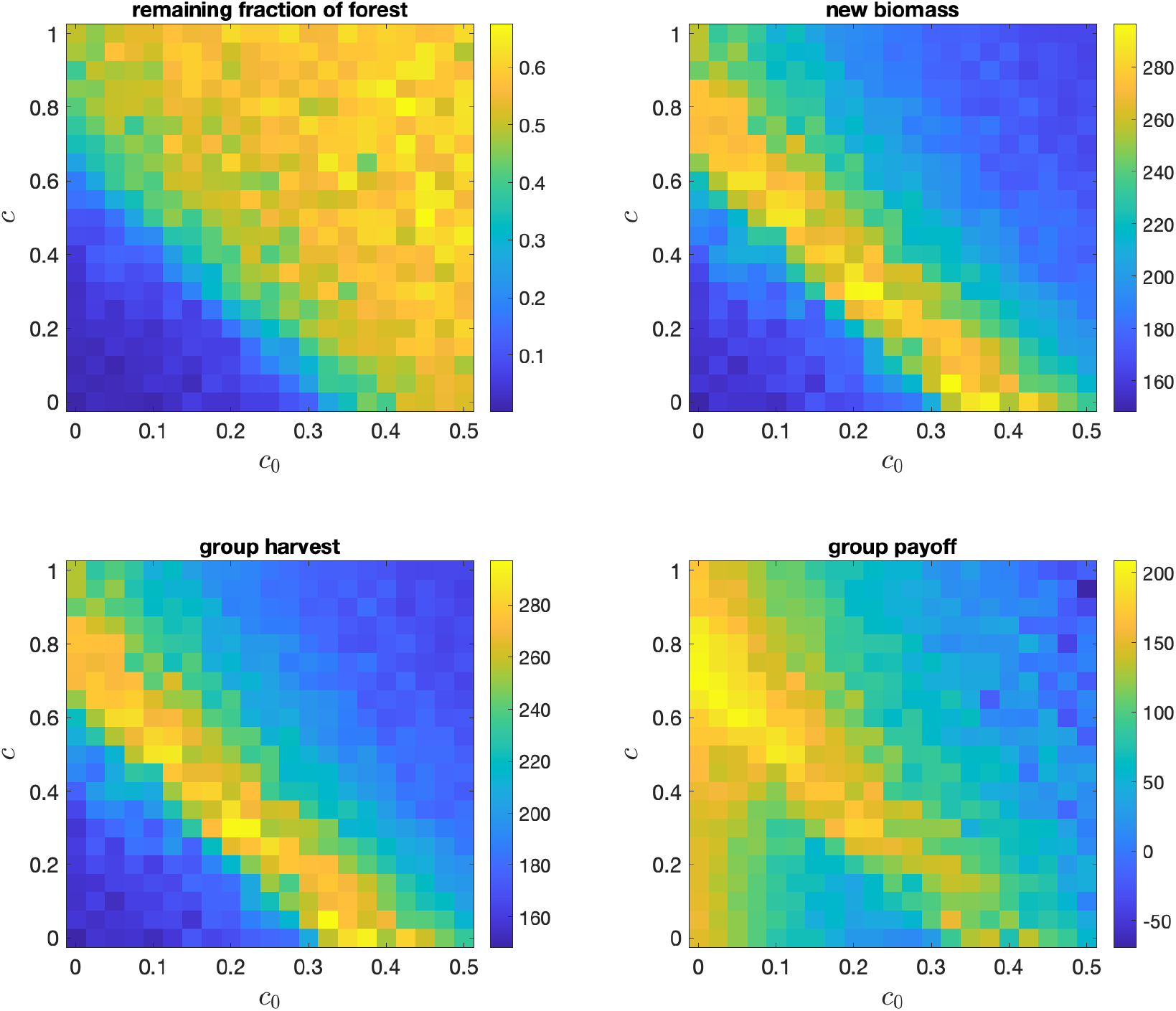
Effects of the cost parameters *c*_0_ and *c* on: the remaining fraction of forest *f*, the new biomass 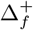, the group harvest *x*_*a*_, and the group payoff *π*. Shown are the averages based on 100 runs each of 1000 rounds for each parameter combination. Results in each run are averages based on the last 250 rounds. *F* = 0.95, other parameters are at default values. **The results for** *F* = 0.5 **are shown in Figure S3**.

**Figure S10:**
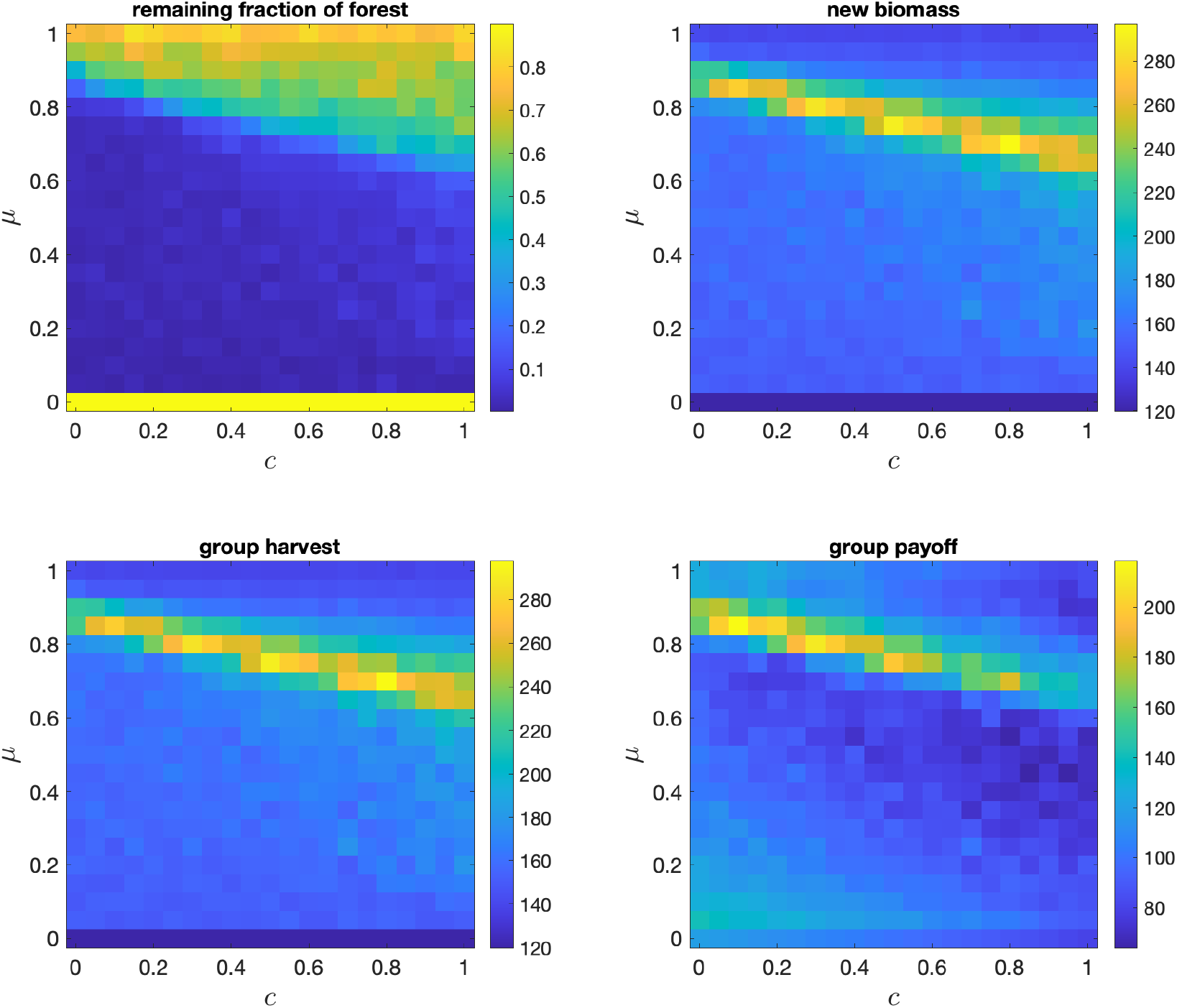
The joint effects of the cost parameter *c* and the relative importance of normative reasoning *µ* on: the remaining fraction of forest *f*, the new biomass 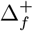, the group harvest *x*_*a*_, and the group payoff *π*. Shown are the averages based on 100 runs each of 1000 rounds for each parameter combination. Results in each run are averages based on the last 250 rounds. *F* = 0.95, other parameters are at default values. **The results for** *F* = 0.5 **are shown in Figure S4**.

**Figure S11:**
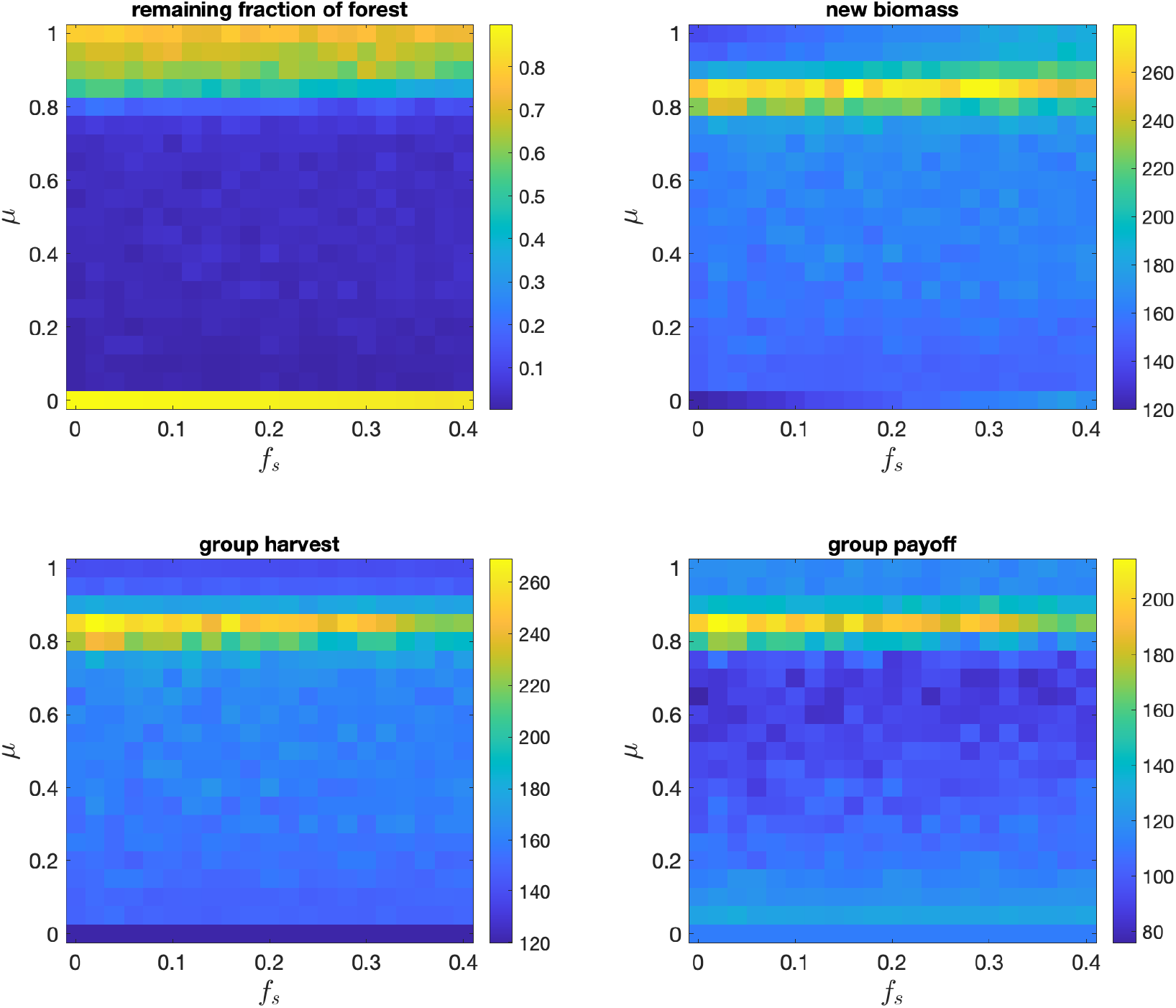
The effects of the relative importance of normative reasoning *µ* and the severity of shocks *f*_*s*_ on: the remaining fraction of forest *f*, the new biomass 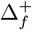, the group harvest *x*_*a*_, and the group payoff *π*. Shown are the averages based on 100 runs each of 1000 rounds for each parameter combination. Results in each run are averages based on the last 250 rounds. Each round, a shock occurs randomly and independently with probability *p*_*sh*_ = 0.01. *F* = 0.95, other parameters are at default values. **The results for** *F* = 0.5 **are shown in Figure S5**.

### S2.2 Supplementary Figures for the case of *F* = 0.95

In this section we redo the entire analysis presented in the main text for the case of *F* = 0.95.

### S2.3 Supplementary Figures for comparison of cases with *F* = 0.5 **and** *F*

**Figure S12:**
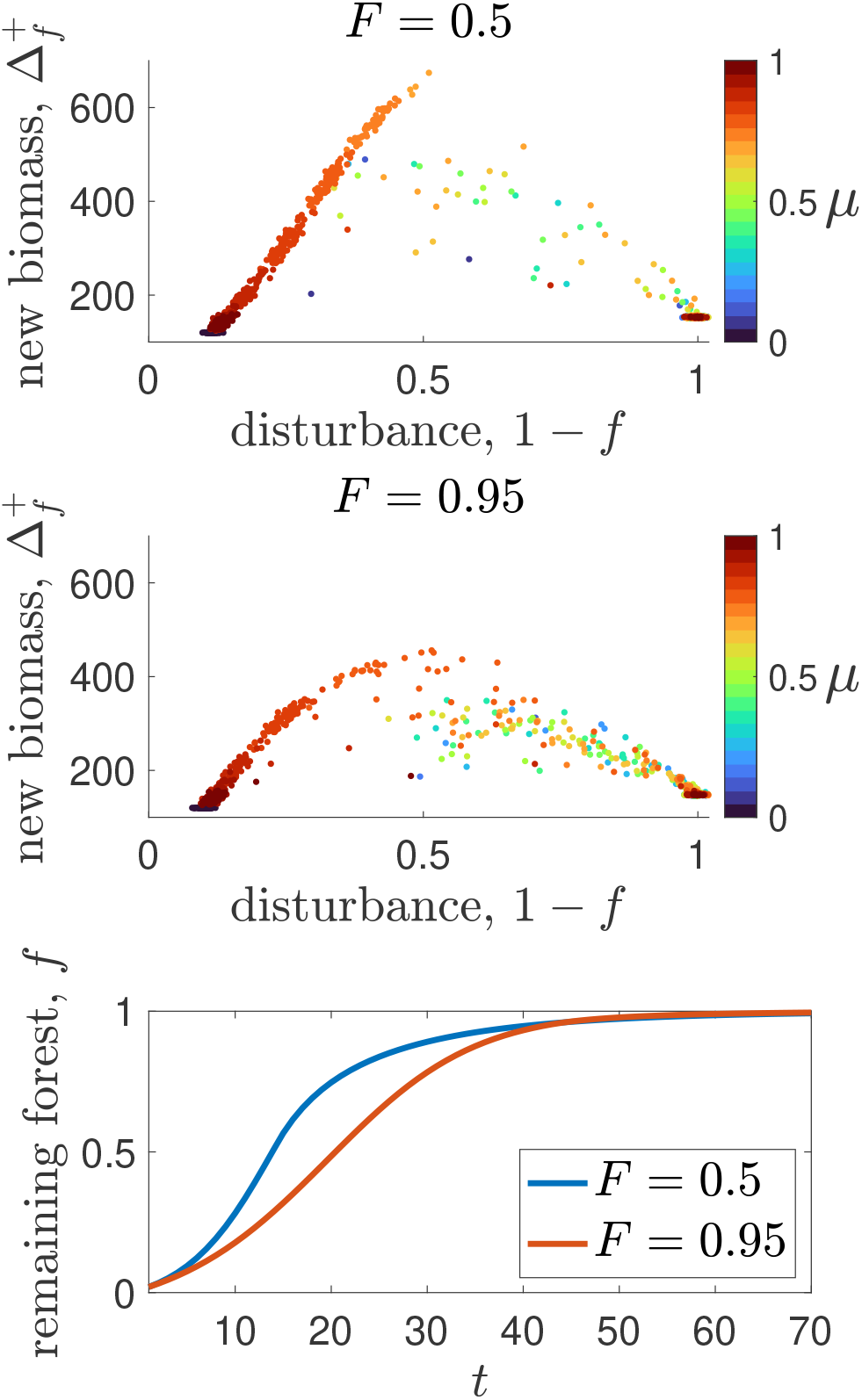
Top and middle panels: the effects of parameter *F* on equilibrium levels of the amount of new biomass 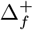 and disturbance (1 − *f*) (the effects on **harvest** 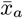 are shown in Figure 5a). Results are shown for *F* = 0.5 and *F* = 0.95. Each point is the average of 100 runs for each value of *µ* ∈ *{*0, 0.05, ‥, 1*}*. The value of each run is an average of the final 250 rounds out of 1,000 total rounds. Other parameters are at default values. Bottom panel: mean-field approximation (see Section S1.2 of the SM) for the reforestation curves in the absence of anthropogenic impact.

## S3 Supplementary code

This section contains Matlab code that reproduces the results of numerical simulations. Main.m performs simulations using the function dis opt .m, which calculates the myopic best response.

### S3.1 Main.m

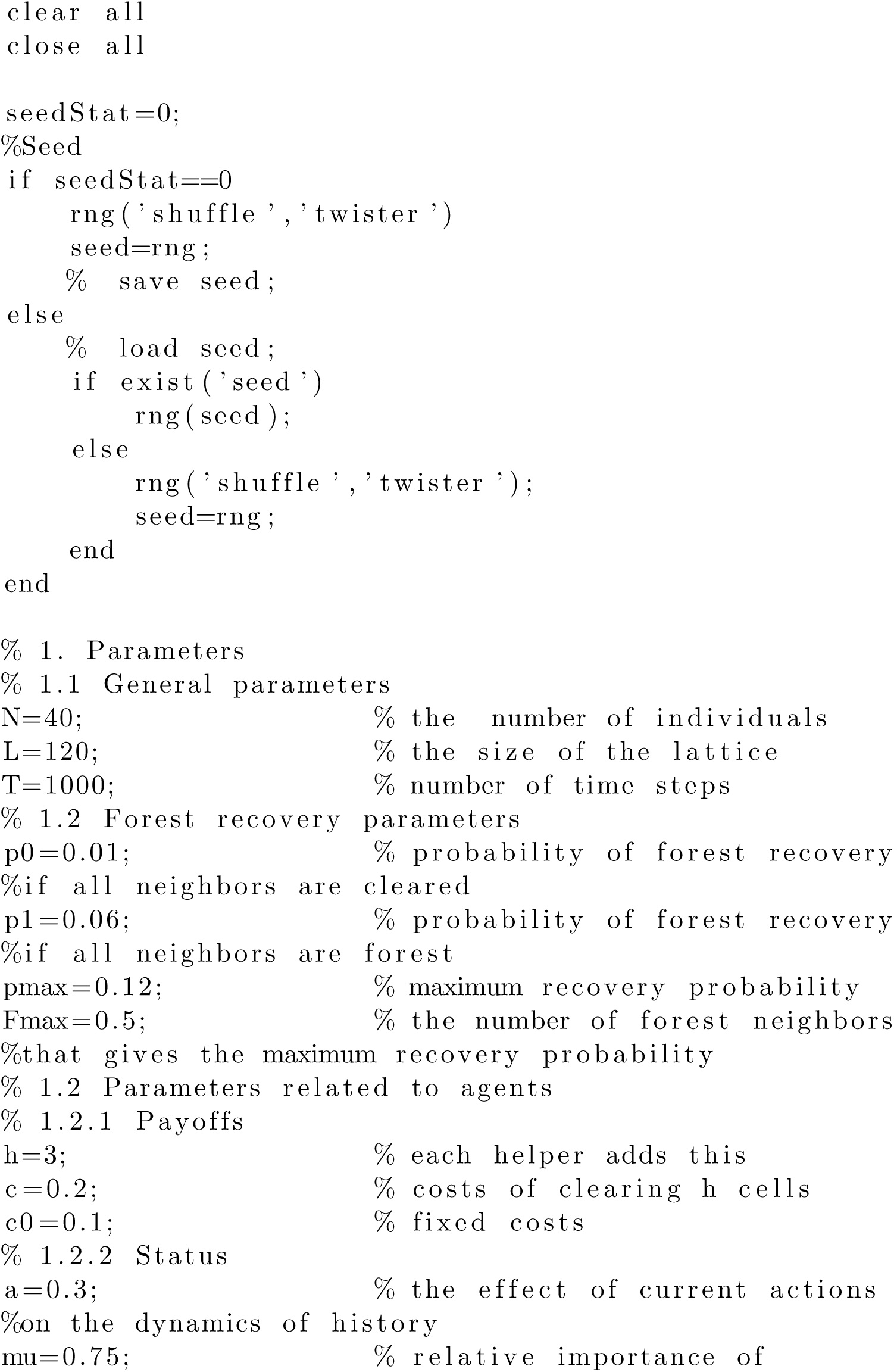
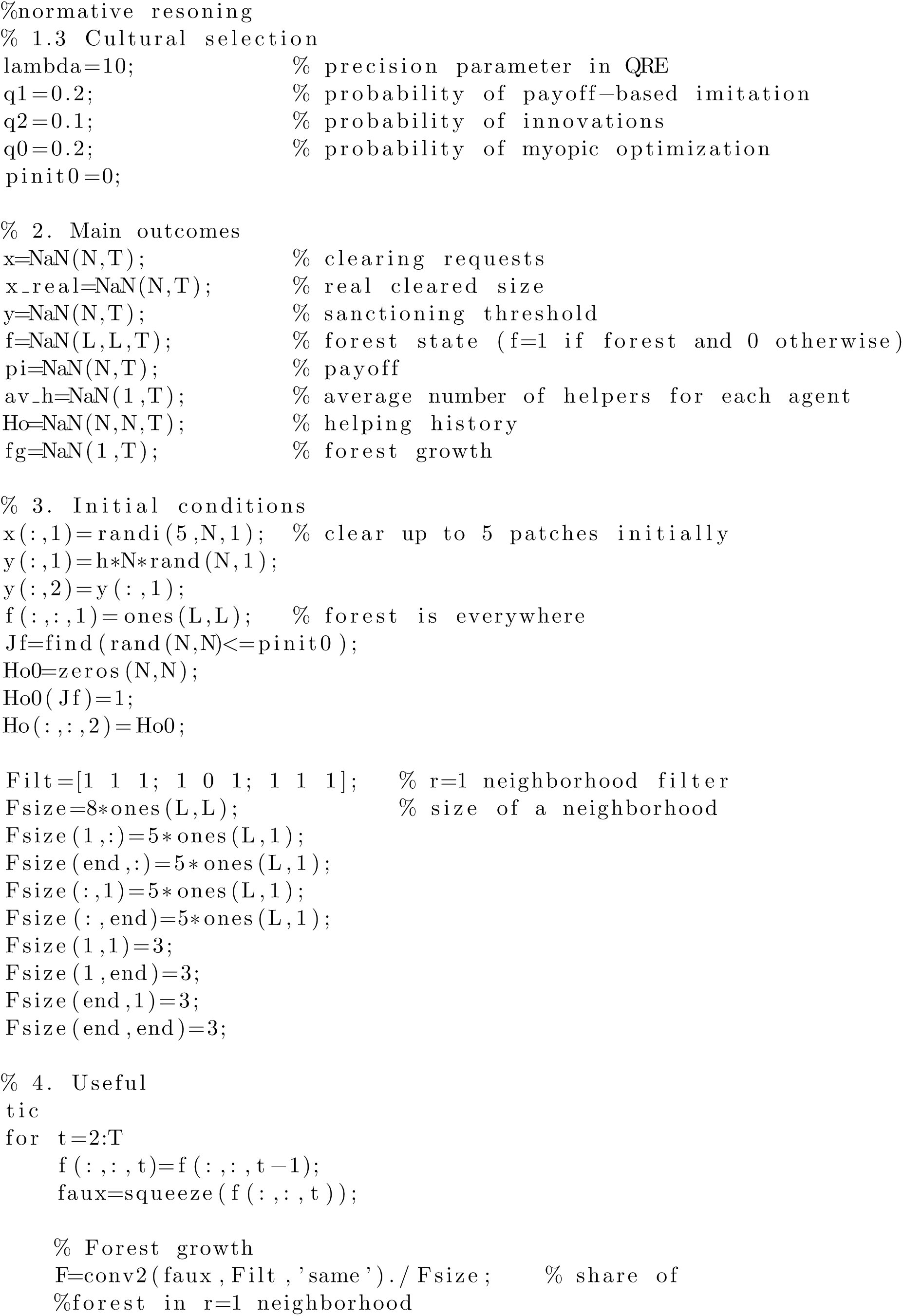
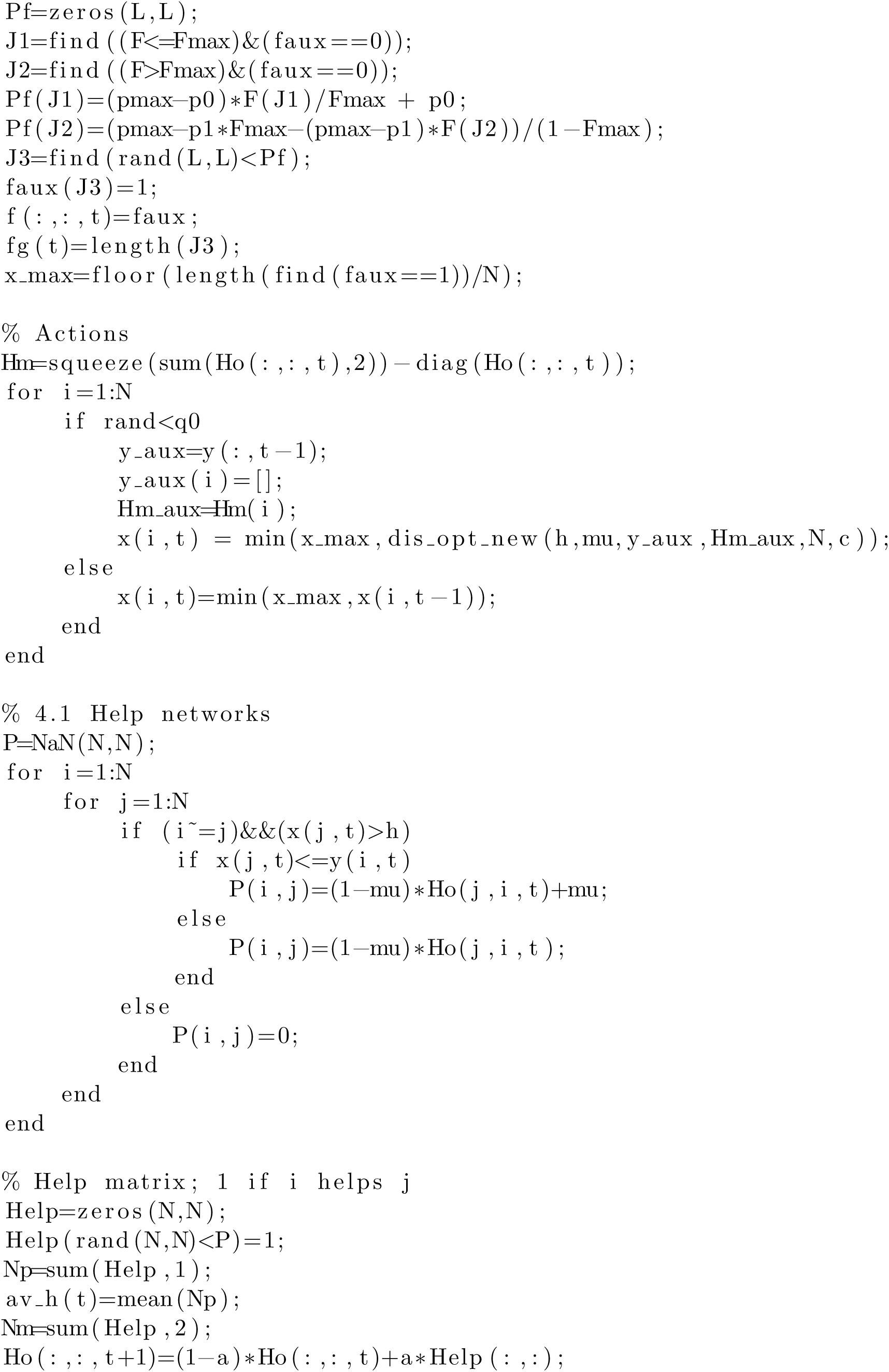
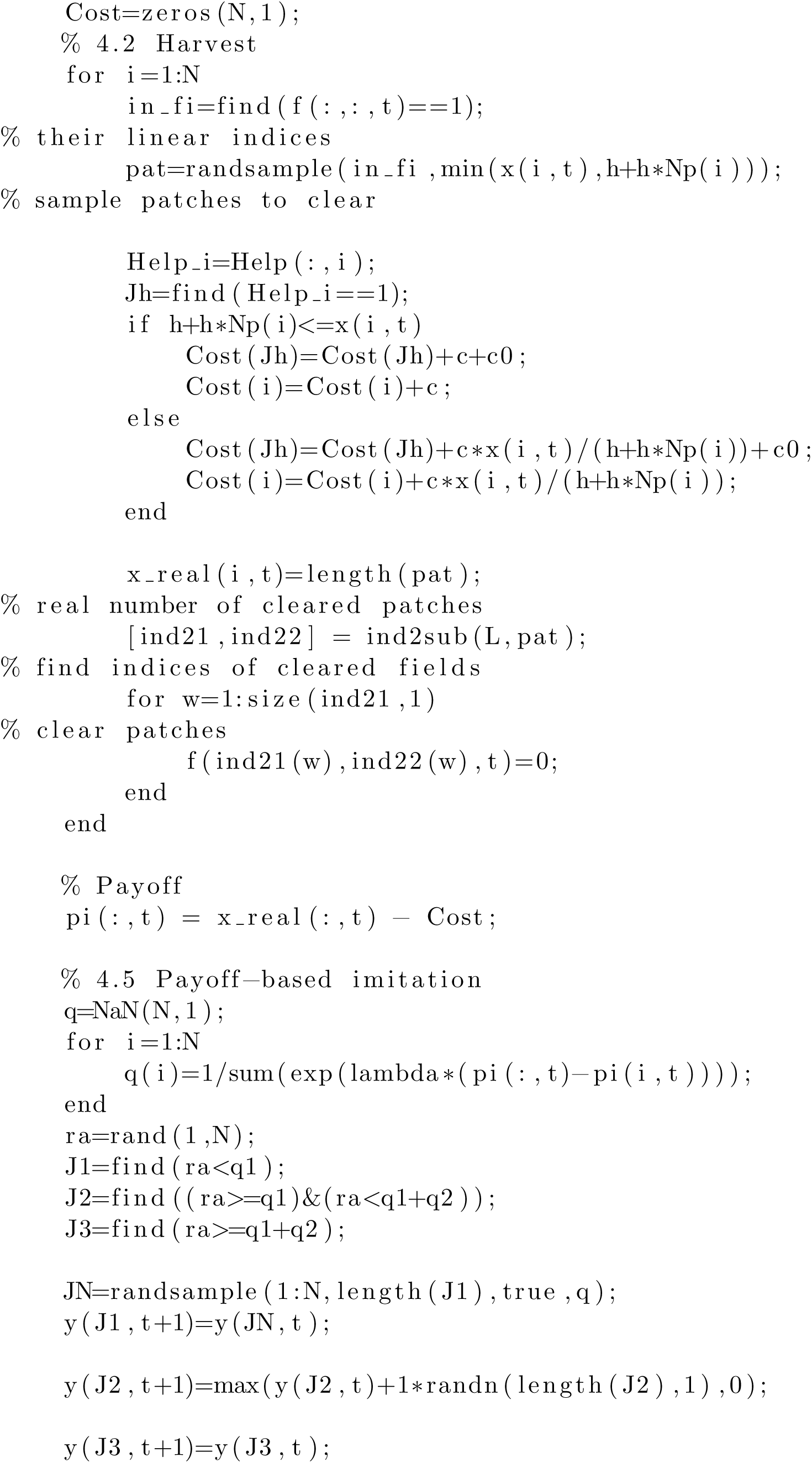
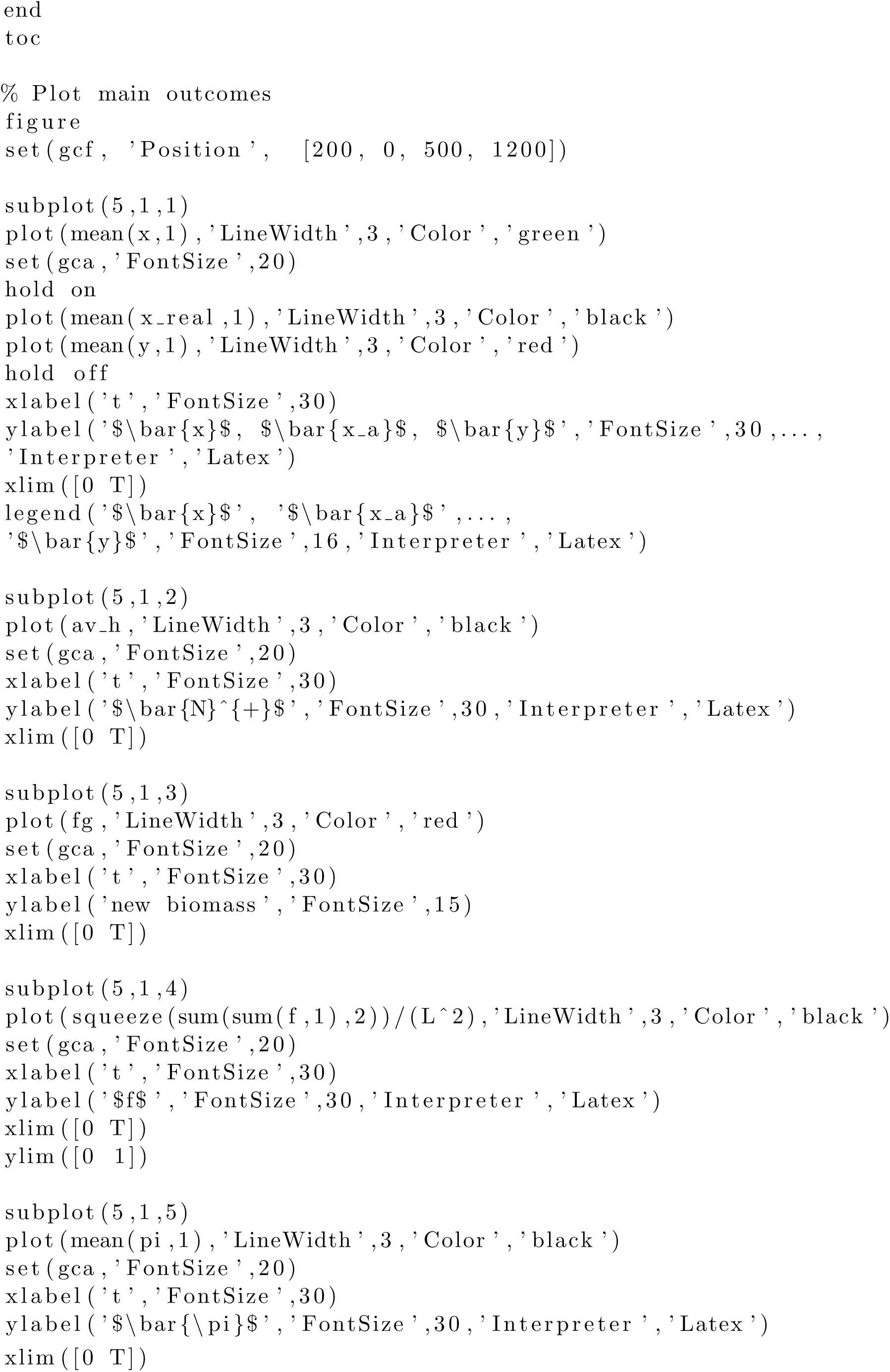

### S3.2 dis_opt_new.m

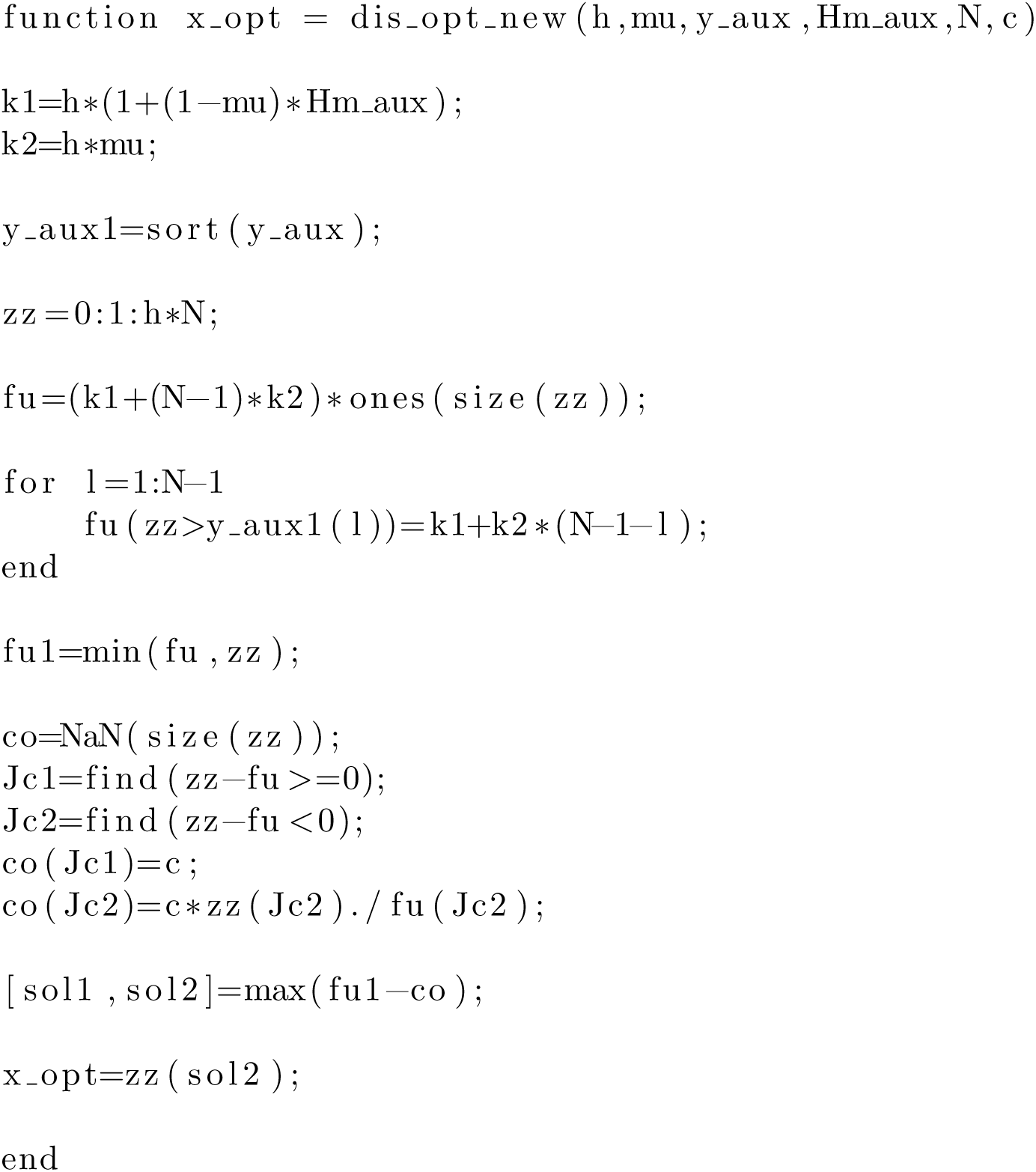

## Footnotes

1 The maximum number of cells each player was allowed to clear each round of the game was five.

2 In this case, the dynamics of the remaining fraction of forest can be approximately described by logistic growth (for more details see Section S1.2 of the SM).

3 We postulate *q*_*i,j*_ = 1 if *i* = *j*.

4 Note this effect only holds if costs of helping are sufficiently high.

5 Note that in sustainable outcomes, *f* is typically greater than 0.5. Some analytical insights explaining this phenomenon can be found in Section S1.2 of the SM.

## Notes

### Competing Interest Statement

The authors have declared no competing interest.

